# Timed receptor tyrosine kinase signaling couples the central and a peripheral circadian clock in *Drosophila*

**DOI:** 10.1101/2023.05.14.540706

**Authors:** Javier Cavieres-Lepe, Emad Amini, Dick R. Nässel, Ralf Stanewsky, Christian Wegener, John Ewer

**Affiliations:** Centro Interdisciplinario de Neurociencias de Valparaíso, Universidad de Valparaíso, Valparaíso, Chile; Julius-Maximilians-Universität Würzburg, Biocenter, Theodor-Boveri-Institute, Neurobiology and Genetics, Am Hubland, 97074 Würzburg, Germany; Department of Zoology, Stockholm University, 10691 Stockholm, Sweden; Institute of Neuro-and Behavioral Biology, Multiscale Imaging Centre, University of Münster, 48149 Münster, Germany; Instituto de Neurociencias de Valparaíso, Universidad de Valparaíso, Valparaíso, Chile

**Keywords:** Keywords: Prothoracic gland, eclosion, PTTH, neuropeptide, circadian rhythms

## Abstract

Circadian clocks impose daily periodicities to behavior, physiology, and metabolism. This control is mediated by a central clock and by peripheral clocks, which are synchronized to provide the organism with a unified time through mechanisms that are not fully understood. Here, we characterized in *Drosophila* the cellular and molecular mechanisms involved in coupling the central clock and the peripheral clock located in the prothoracic gland (PG), which together control the circadian rhythm of emergence of adult flies. The time signal from central clock neurons is transmitted via small neuropeptide F (sNPF) to neurons that produce the neuropeptide Prothoracicotropic Hormone (PTTH), which is then translated into circadian oscillations of Ca^2+^ concentration and daily changes in PTTH levels. Rhythmic PTTH signaling is required at the end of metamorphosis, and transmits time information to the PG by imposing a daily rhythm to the expression of the PTTH receptor tyrosine kinase (RTK), TORSO, and of ERK phosphorylation, a key component of PTTH transduction. In addition to PTTH, we demonstrate that signaling mediated by other RTKs contribute to the rhythmicity of emergence. Interestingly, the ligand to one of these receptors (Pvf2), plays an autocrine role in the PG, which may explain why both central brain and PG clocks are required for the circadian gating of emergence. Our findings show that the coupling between the central and the PG clock is unexpectedly complex and involves several RTKs that act in concert, and could serve as a paradigm to understand how circadian clocks are coordinated.

**Significance statement:** Circadian clocks impose daily periodicities to behavior, physiology, and metabolism, and are synchronized to provide the organism with a unified time through mechanisms that are poorly understood. In holometabolous insects, the circadian control of adult emergence depends on the coupling between the central clock and a peripheral clock located in the prothoracic gland (PG). Here we identify the cellular and molecular mechanism that transmits time information from the central clock to the PG clock. This process is unexpectedly complex and involves a number of receptor tyrosine kinases (RTKs). Such a mechanism may add robustness to the coupling between the 2 clocks and serve as a paradigm for understanding how circadian clocks are coordinated.

## Main text

### Introduction

Circadian rhythms allow multicellular organisms to anticipate daily changes in the environment such as the arrival of dawn or dusk. In animals, these behavioral and physiological rhythms are generated by multi-oscillator systems composed of a central pacemaker housed in the brain as well as of peripheral pacemakers located in a wide variety of tissues. The coordination of these clocks is critical for the organism to express a unified circadian time (Mohawk et al., 2012). In mammals, the so-called “master pacemaker” located in the suprachiasmatic nucleus (SCN) synchronizes to the environmental day-night cycles and coordinates peripheral oscillators through neural, endocrine, behavioral, and thermal signals (Welsh et al., 2010). In turn, a variety of peripheral signals feedback to adjust and stabilize SCN rhythmicity (Buijs et al., 2016; Harder and Oster, 2020). However, our understanding of the cellular and molecular mechanisms and rules that mediate this coupling is still fragmentary (Pilorz et al., 2018). In the fruit fly, *Drosophila melanogaster,* the central clock is comprised of about 150 neurons that are critical for imposing a daily periodicity to behaviors such as locomotor activity and sleep (Nitabach and Taghert, 2008). A key component of the central clock are the small and large ventral lateral neurons (s-and lLNvs, respectively), which produce the neuropeptide pigment-dispersing factor (PDF) and are critical for circadian timekeeping (Shafer and Yao, 2014). In addition, peripheral clocks reside in a variety of tissues and are mostly autonomous and entrained directly by external inputs (Ivanchenko et al., 2001; Glaser and Stanewsky, 2005; Ito and Tomioka, 2016). However, in other cases, and similarly to mammals, the central clock can coordinate its activity with peripheral oscillators. In particular, the rhythm of emergence of adult flies (used here interchangeably with the term, eclosion), is mediated by the coupling of the brain clock and a peripheral oscillator that resides in the prothoracic gland (PG), the endocrine gland that produces the steroid molting hormone, ecdysone (Myers et al., 2003; Morioka et al., 2012; Selcho et al., 2017). We previously reported that this coupling is mediated by a peptidergic signaling pathway in which sNPF from the sLNv clock neurons inhibits the neurons that express PTTH (Selcho et al., 2017). In turn, the PTTH neurons (PTTHn) secrete PTTH, which binds to the receptor tyrosine kinase (RTK) encoded by *torso* in cells of the PG, to control the biosynthesis of the molting hormone, ecdysone (McBrayer et al., 2007; Selcho et al., 2017). Although the titers of this steroid must fall below a threshold level to trigger emergence (Truman, 1984; Porter and Collins, 2009), we recently showed that the clock does not exert its action by regulating the levels of ecdysone but by controlling its actions, which are mediated by the ecdysone receptor in the PG (Mark et al., 2021). Indeed, although injections of 20E delay the time of eclosion they do not affect its circadian gating. By contrast, disabling 20E actions in the PG renders arrhythmic the pattern of adult emergence.

Although the neuropeptide circuit that connects the brain clock to the PG clock has been identified, how and when during development the time signal is transmitted from the central circadian pacemaker to this peripheral clock remains unclear. For example, although sNPF from sLNvs suppresses Ca^2+^ activity in the PTTHn (Selcho et al., 2017), it is not known whether this translates into a rhythmic activity and output from PTTHn. Similarly, although PTTH is required for the circadian rhythmicity of emergence, *ptth* mRNA levels do not exhibit a circadian fluctuation (Selcho et al., 2017), indicating that timing information from the central clock may be encoded by a different mechanism. In addition, PTTHn are targets of peptidergic inputs other than sNPF at least in larvae (Deveci et al., 2019; Imura et al., 2020; Hao et al., 2021), yet it is not known whether any of these peptides also affect the circadian rhythm of emergence. Finally, signaling molecules other than PTTH, including Jelly belly, PDGF-and VEGF-related factor, Egf, and insulins, also regulate ecdysone production by the PG (Colombani et al., 2005; Cruz et al., 2020; Pan and O’Connor, 2021) but their role in the circadian control of the eclosion is currently unknown.

In order to understand how and when the time signal is transmitted from the central clock to the PG clock, we examined the anatomical, cellular, and molecular basis of the central clock-PTTHn-PG axis. We first determined the connectivity between sLNv clock neurons and the PTTHn by pre-and postsynaptic tracing, and found that this connection is unidirectional and likely to be exclusively peptidergic. We also show that PTTHn exhibit a daily oscillation in intracellular Ca^2+^ levels ([Ca^2+^]i) and of PTTH immunoreactivity at the terminals of PTTHn in the PG. Unexpectedly, we found that PTTH is required for the circadian rhythmicity of *torso* expression and of PTTH transduction in the PG. Direct imaging of the activity of PTTHn and the PG revealed that the final peak of activity in PTTHn occurred around 16h before eclosion and was followed 6h later by a peak of activity in the PG . This timing is consistent with our recent findings on the events that control the timing of eclosion (Mark et al., 2021), and we further show here that PTTHn signaling is required at the end of metamorphosis for the expression of a circadian rhythm of adult emergence. In addition, the presence of activity peaks in the PG that occurred prior to the activation of PTTHn suggests that the PG responds to additional factors, and we show here that PDGF-and VEGF-receptor related (Pvr) and Anaplastic lymphoma kinase (Alk), two RTKs that are also expressed in the PG, contribute to the circadian control of emergence. Thus, our detailed characterization of the transduction pathway from the sLNv clock neurons to the PG clock reveals that the transmission of time information between clocks is surprisingly complex and involves multiple actors. It also defines a new role for RTK signaling in the coordination of circadian clocks. Our work may provide a general mechanism for understanding how circadian clocks are coordinated.

## Results

### All PDF-positive sLNv signal non-synaptically to the PTTHn

The dorsal projections of PDF-expressing LNv clock neurons terminate in close proximity to PTTHn arborizations (Fig. 1A-A”). We previously showed that sLNvs transmit time information to PTTHn via peptidergic sNPF signaling, but not PDF(Selcho et al., 2017). To determine which LNv neurons signal to PTTHn and whether this contact includes synaptic transmission, we performed a connectomic analysis by GFP reconstitution across synapses (syb-GRASP) (Macpherson et al., 2015), BAcTrace (Cachero et al., 2020), and *trans-Tango* MkII (Sorkac et al., 2022), in adult pharate animals. This work was necessary because the PTTHn can no longer be detected starting shortly after eclosion (Liu et al., 2016) and are therefore not included in the available EM-based connectomic datasets that are based on older adult flies (Scheffer et al., 2020). Despite the close proximity of the sLNv terminals to the dendritic arborizations of PTTHn, the reconstituted GFP signal between these neurons using syb-GRASP was weak and restricted to the primary process (Figure 1B), suggesting that sLNv transmit time information to the PTTHn primarily via non-synaptic connections. We then used BAcTrace and *trans-Tango MkII*, which are retrograde and anterograde neuronal tracers, respectively, to determine the directionality of the connection, and to identify the LNvs that provide input to the PTTHn. As shown in Figures 1C and 1D, BAcTrace labeled all sLNv and none of the lLNv, whereas *trans-Tango MkII* labeled both pairs of PTTHn. Finally, we used receptor-specific intragenic driver lines to show that PTTHn express the receptor for sNPF (sNPFR) but not that for PDF (PDFR) (Fig. 1E and 1F), consistent with our previous finding that transmission is mediated via sNPF and not PDF (Selcho et al., 2017).

**Figure 1.**
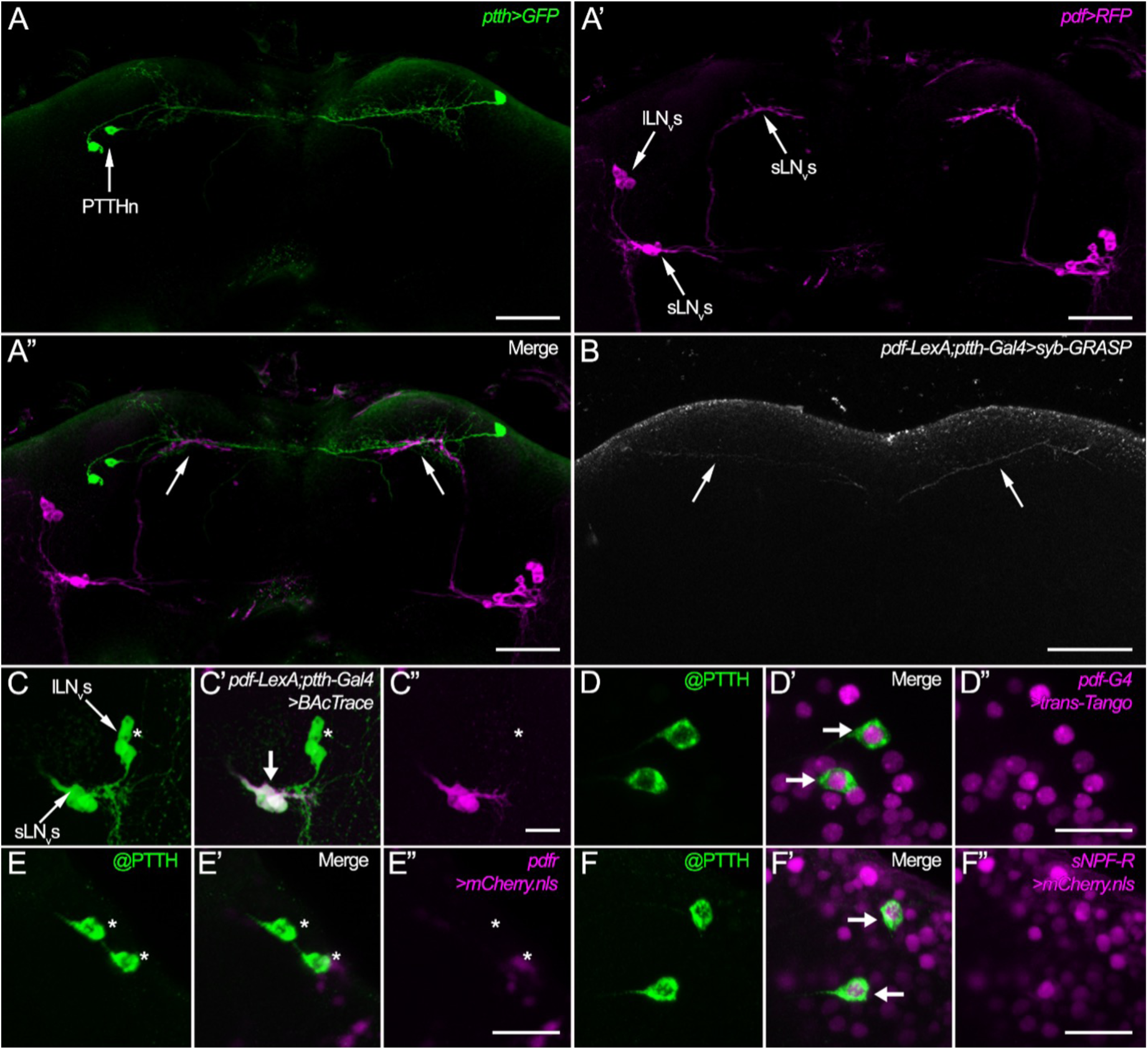
Connectivity between PDF-expressing sLNv clock neurons and PTTHn. **(A-A”)** The arborizations of PTTHn in the superior protocerebrum **(A)** are in close contact (arrows in **(A”)**) with the PDF-expressing sLNv clock neurons **(A’)**. **(B)** *syb*-GRASP reconstruction between PTTHn and PDF-expressing sLNv clock neurons produced only a faint reconstruction of GFP (arrows), which did not include small branches in the contact area between PTTHn and sLNv located in the superior protocerebral region. **(C-C”)** BAcTrace-labeling showing that all PDF-expressing sLNv but not the large lLNv synapse onto the PTTHn. Whereas *pdf*-LexA drives GFP expression in the sLNv and lLNv **(C)**, the BAcTrace signal **(C’)** was only detected in the sLNv **(C”)**. **(D-D”)** *Trans-Tango* MkII labeling showed that all PTTHn are downstream to the sLNv since the positive *trans-Tango* MkII signal was visible in PTTHn present in each hemisphere **(D’)**. **(E-E”).** A PDF receptor-specific intragenic driver line **(E”)** did not label the PTTHn **(E)** in pharate adults, indicating that PDF receptor is not expressed **(E’)**. Asterisks mark the position of the PTTHn. **(F-F”).** By contrast, an sNPF receptor-specific intragenic driver line **(F”)**, labeled the PTTHn **(F)** in pharate adults **(F’),** indicating sNPF receptor expression. Scale bars: A-B: 50 *μ*m; C-F: 20 *μ*m.

### Ca^2+^ signaling in PTTHn is relevant to the circadian control of adult emergence

Since sNPF released from sLNv reduces Ca^+2^ levels [Ca^2+^] in PTTHn, we explored whether [Ca^2+^] in PTTHn neurons changed in a time-dependent manner. For this we used the genetically encoded Ca^+2^ sensor, GCaMP6M (Chen et al., 2013), to measure changes in [Ca^+2^] in PTTHn during the course of the day (and subjective day) at the beginning of metamorphosis. (White pre-pupal stage [WPP] animals were used because the circadian clock is intact and fully functional at this time (Morioka et al., 2012). By contrast, the PG of animals close to emergence is undergoing apoptosis (Dai and Gilbert, 1991), making them difficult to image.) As shown in Figure 2A-C, the cell bodies of PTTHn showed a daily rhythm in [Ca^+2^] under a 12h light: 12h dark regime (LD), with maxima at the beginning of the night (Zeitgeber time (ZT) 12) and minima 4h after lights-on (ZT4). This daily oscillation was maintained under constant darkness (DD), but the peak was delayed by about 6h (circadian time (CT) 18; where CT12 is the start of the subjective night) (Figure 2E) (this delay may occur because photic inputs can set the Ca^+2^ phase in a group of clock neurons (Liang et al., 2017)). Importantly, this daily rhythm was lost under both LD (Figure 2D) and DD conditions (Figure 2F) in animals bearing null alleles for the *period* gene (*per^0^*), which is a core component of the clock. Expression of GCaMP in the PTTHn did not affect the rhythm or period of emergence (Table S1).

**Figure 2.**
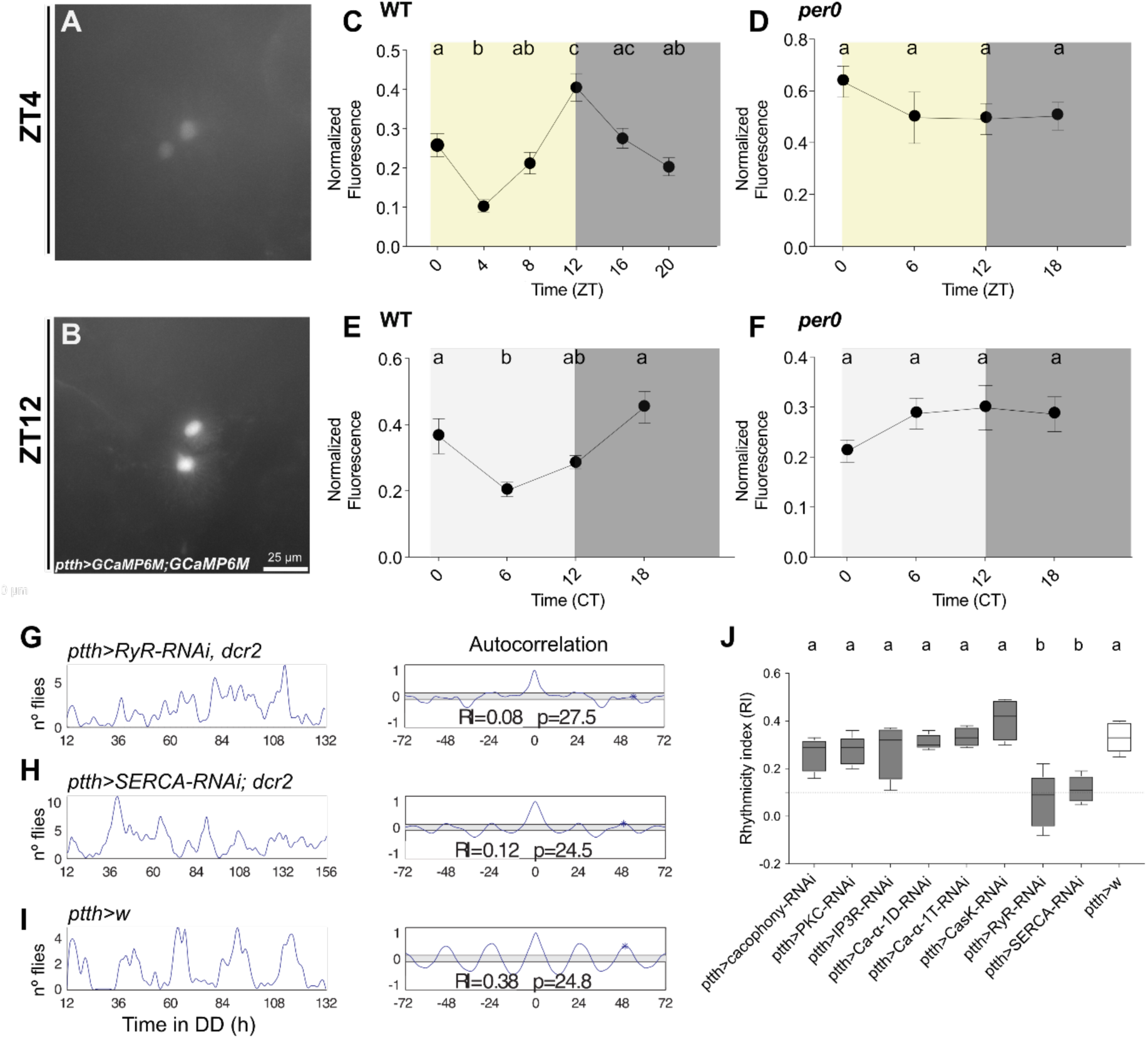
Ca^+2^ signaling in PTTHn is relevant to the circadian control of adult emergence. (A-B) Representative images of GCaMP fluorescence in PTTHn in the brains of WPP examined at ZT4 (A) and ZT12 (B). (C-F) Levels of GCaMP fluorescence in PTTHn of: wildtype (WT) animals at different times of day under LD (C) and DD conditions (E), and in the soma of PTTHn of *per[0]* mutants under LD (D) and DD (F) conditions. Different letters indicate statistically different groups. (ANOVA with Tukey’s *post hoc* test). Between 8 and 10 animals were analyzed for each timepoint. ZT: zeitgeber time. CT: Circadian time. (G-I) Records showing the time course of emergence of a single population of flies under DD conditions (left) and corresponding autocorrelation analysis (right) of populations bearing a knockdown in PTTHn of Ryanodine Receptor (RyR)(G), SERCA (H), and in corresponding controls. (I). Periodicity (p, in hours) and associated rhythmicity index (RI) are indicated. (J) Average RI values from knockdown in PTTHn of mediators of Ca^+2^ signaling. The dashed line at RI=0.1 indicates cutoff below which records are considered arrhythmic. Different letters indicate statistically different groups. (p<0.05; one-way ANOVA, Tukey’s *post hoc* multiple comparison analyses).

In order to determine if there is a temporal relationship between the rhythm of sNPF release from sLNv and the Ca^+2^ oscillations in PTTHn, we measured the temporal pattern of neuropeptide abundance at the terminals of sLNv by expressing ANF-GFP, a transgenic neuropeptide reporter (Husain and Ewer, 2004), in PDF-expressing LNv. As shown in Figure S1A-S1B, and similar to previous findings for PDF in adult flies(Park et al., 2000), the ANF-GFP signal in the terminals of sLNv at the WPP stage was highest at the beginning of the day, which is coincident with the time when [Ca^+2^] levels in PTTHn were minimal (Figure 2C). Thus, our results suggest that the phase of the Ca^+2^ rhythm in PTTH neurons is set by the inhibitory sNPF input from sLNv (although *pdf*>ANF-GFP reports the abundance of both sNPF and PDF, PTTHn do not express PDFr).

In order to evaluate the relevance of Ca^+2^ in the PTTHn to the circadian control of adult emergence, we knocked-down elements involved in Ca^+2^ signaling in PTTHn and determined the consequences on the rhythm of adult emergence (Figure 2G-2J). We found that the knockdown of voltage-gated Ca^+2^ channels (Ca^+2^-α-1D, Ca^+2^-α-1T, and *cacophony*) and of Ca^+2^-binding proteins (CasK, and PKC) did not affect the rhythm or the periodicity of eclosion (Figure 2J). By contrast, knocking down the expression of RyR (ryanodine receptor), an endoplasmic reticulum Ca^+2^ channel, caused a significant weakening of the circadian pattern of emergence (Figure 2G and 2J). (In these and all RNAi-mediated knockdown experiments, at least 2 different RNAi transgenes were tested; see Materials and Methods.) A similar result was obtained following the knockdown of SERCA (sarco/endoplasmic reticulum calcium ATPase) (Figures 2H and 2J). Importantly, knockdown of RyR or SERCA did not affect the gross morphology of PTTHn (Figure S2A-S2C) suggesting that the weakening of the rhythm of the emergence may be caused by alterations in Ca^+2^ homeostasis in PTTHn. Together, these results reveal that Ca^+2^ signaling in PTTHn through the RyR and SERCA transporters is part of the cellular mechanism that imposes a daily rhythm to the pattern of adult emergence

### The PTTH/torso axis is under circadian control

Although PTTH transmits time information from the sLNv to the PG clock, this timing signal does not involve the circadian regulation of *ptth* transcript levels (Selcho et al., 2017). However, whether PTTH abundance cycles in the PTTHn axon terminations in the PG is unknown. To explore this possibility, brain-PG complexes from WPP were dissected at different times of day and immunostained for PTTH. As shown in Figure S3A, PTTH immunosignal showed a rhythm in the cell bodies; however we found that this rhythm was not dependent on the clock. (This cycling of PTTH staining in the cell bodies may be dependent on photic inputs, similar to the role of PTTH in light avoidance during larval stages (Yamanaka et al., 2013; Sorkac et al., 2022).) By contrast, PTTH immunoreactivity of the axonal terminals onto the PG showed clock-dependent daily rhythmicity. PTTH immunoreactivity was maximal during the night and early morning and minimal during the day (Figure 3A-3C), persisted under DD conditions, and was abolished in *per^0^* null mutant animals under both LD and DD conditions (Figure 3D-3F). Together, these results indicate that the circadian clock acts at the post-translational level to impose a daily rhythm of PTTH accumulation at the site of innervation of the PG and suggest that the PTTH neuropeptide may be rhythmically released onto this gland.

**Figure 3.**
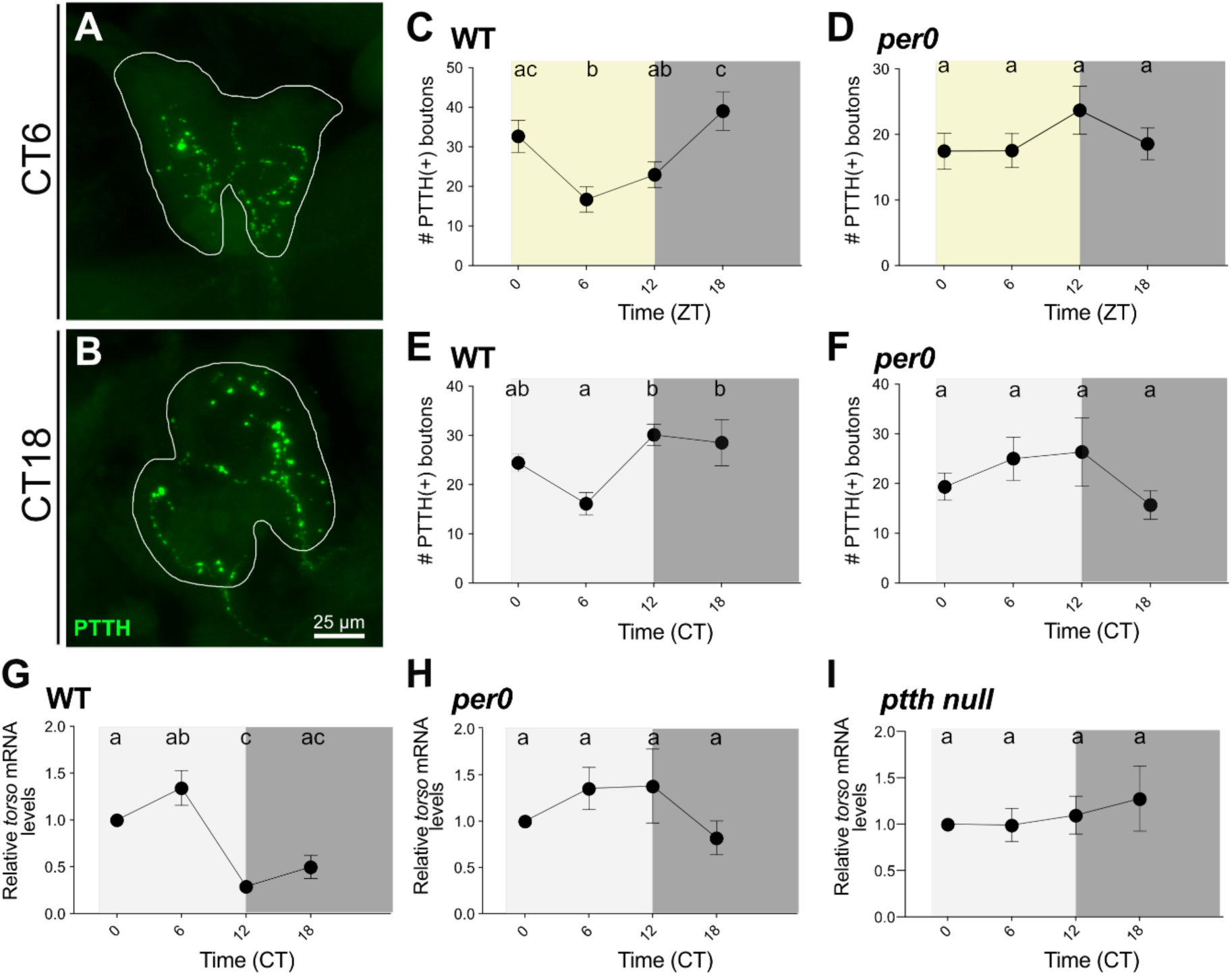
Circadian regulation of PTTH/TORSO axis. (A-B) Pattern of PTTH-immunoreactivity of terminals of PTTHn on the PG of WPP animals at CT6 (A) and CT18 (B). White line delimits the PG. (C-F) Average number of PTTH-immunoreactive boutons showing above-threshold intensity on the PG at different times of day in wildtype (C, E) and in *per[0]* mutants (D,F) under LD (C,D) and DD (E,F) conditions. Eight–10 brains were analyzed per time-point. (G-I) Relative expression of mRNA levels of *torso* in PGs isolated from WPP stage wildtype (G), *per[0]* (H), and *ptth* null mutant (I) animals under DD conditions. Different letters indicate statistically different groups (p<0.05; one-way ANOVA, Tukey’s *post hoc* multiple comparison analyses).

Another critical step in the transmission of time information from the brain clock to the PG clock is the PTTH receptor, TORSO (Selcho et al., 2017). To test whether *torso* expression is regulated by the clock, we first examined *torso* mRNA abundance in the PG at different times of day, in wildtype and in *per^0^* null mutant animals. As shown in Figure 3G-3H and S3B, *torso* transcript levels exhibited a daily rhythm under LD and DD conditions in wildtype but not in *per^0^* null animals. Intriguingly, this oscillation peaked during the day/subjective day and reached a minimum at the beginning of the night/subjective night, which is in antiphase relative to the oscillations observed for the PTTH-immunoreactivity of the terminals of PTTHn onto the PG. Interestingly, *torso* mRNA levels did not cycle in the PG of *ptth* null mutant animals under DD condition (Figure 3I). Taken together our findings suggest that the circadian rhythmicity of *torso* transcript abundance is dependent on the central clock input, and that this timing signal is transmitted by PTTH. As expected, given that knockdown of *torso* in the PG causes the expression of an arrhythmic eclosion pattern (Selcho et al., 2017), *ptth* null mutant animals showed an arrhythmic pattern of emergence (Figure S3C and S3D).

### Circadian regulation of PTTH transduction in the PG

Our findings reveal that PTTH immunoreactivity of PTTHn terminals and *torso* transcript levels oscillate in antiphase. In order to obtain a readout of the temporal pattern of the resulting activation of the Torso pathway we asked whether the levels of phosphorylated ERK (phosphoERK), a key component of PTTH transduction (Rewitz et al., 2009), express a daily rhythmicity. For this we expressed in the PG a genetically-encoded ERK kinase reporter (ERK-SPARK), which is based on phase separation-based kinase activity (Zhang et al., 2018)(Figure S4A). As shown in Fig. 4, ERK activity was highest during the early part of the day under LD conditions (Fig. 4F); it was also highest during the early part of the subjective day (CT0-2) under DD conditions (Figure 4A, 4B and 4H). Importantly, this daily rhythm in phosphoERK levels was lost in *per^0^* null mutant animals (Figure 4G) indicating that ERK activity in the PG cells is dependent on the circadian clock. This rhythm was also lost when *torso* was knocked down in the PG (Figure 4C, 4D and 4I) (Although rhythmicity itself is lost, a SPARK signal was still detected, as would be expected because this pathway can be activated by other ligands, e.g., insulins). Since early in the day, PTTH immunoreactivity in the PTTHn was lowest and *torso* transcript levels and phosphoERK levels in the PG were highest, our results suggest that PTTH release and Torso activation occur at this time of day. Expression of this reporter in the PG did not affect the circadian rhythmicity of emergence (Figure S4B-S4D).

**Figure 4.**
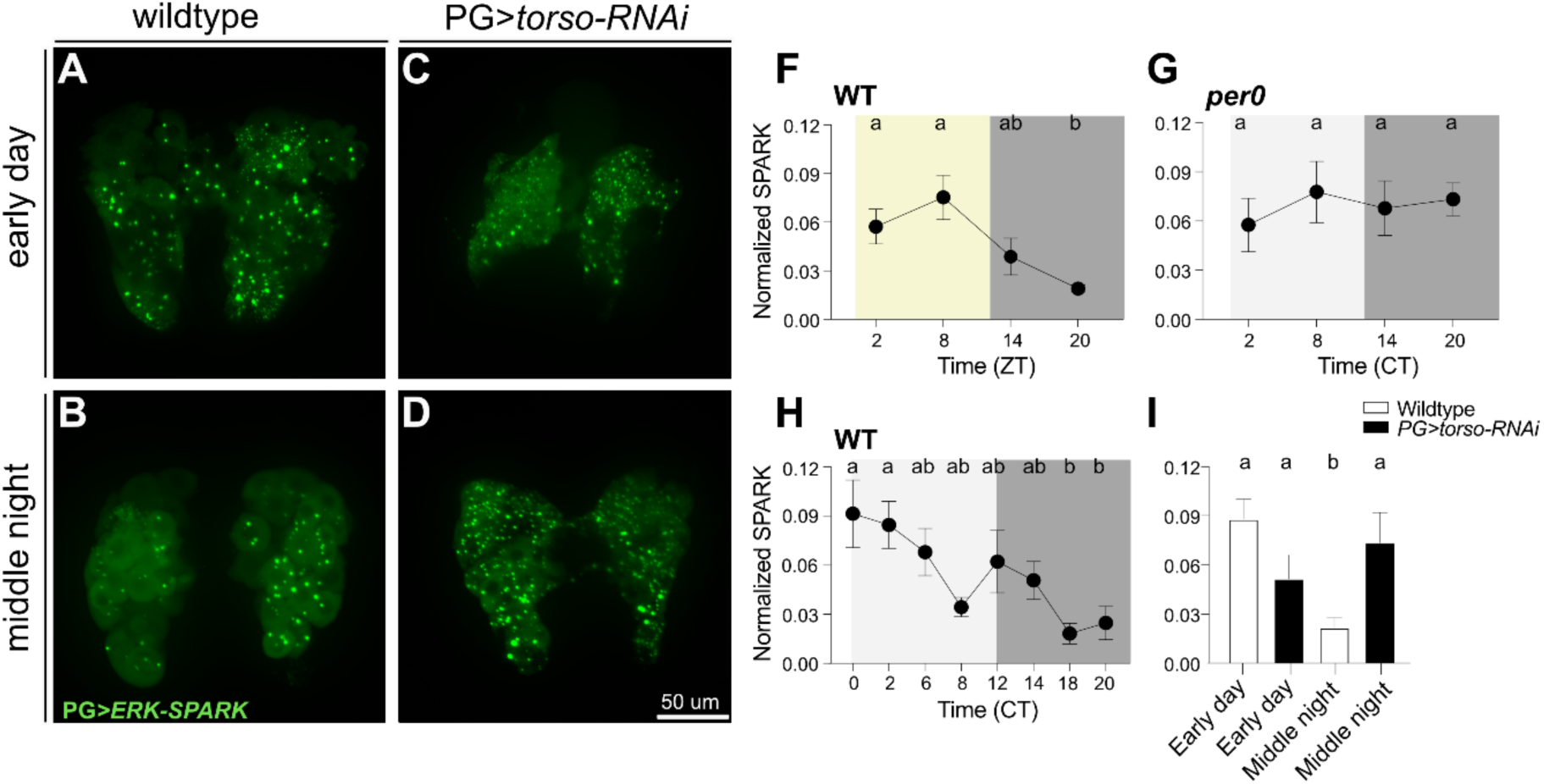
ERK phosphorylation in the PG shows a PTTH-dependent daily rhythm. (A-D) ERK-SPARK signal in whole mount PGs (*phm>ERK-SPARK*) of wildtype (WT) and PG*>torso-RNAi* WPP animals at CT0-2 (early subjective day) and CT18-20 (middle of subjective night). (F-H) Average of normalized ERK-SPARK signal in the PG at different times of day in wildtype PG under LD (F) and DD (H) conditions, and in *per[0]* mutants (under DD condition; G). (I) Average normalized ERK-SPARK signal in wildtype and PG>*torso-RNAi* animals at CT0-2 (early subjective day) and CT18-20 (middle of subjective night). Eight to 10 brains were analyzed per time-point. Different letters indicate statistically different groups (p<0.05; one-way ANOVA, Tukey’s post hoc multiple comparison analyses).

ERK shuttling between cytoplasm and nucleus is relevant for ecdysone biosynthesis and for the control of developmental timing during the larval stages (Ou et al., 2011). As shown in Fig. S5C, nuclear SPARK signal was higher during the day than during the night. This daily oscillation persisted under DD conditions, although the maximum levels were phase-advanced (Figure S5A, S5B and S5BE), which is consistent with the rhythm of the total SPARK signal in the PG cells. In addition, *per^0^* and PG>*torso-*RNAi animals showed an arrhythmic pattern of nuclear SPARK signal (Figure S5D and S5F) indicating that the daily change in phosphoERK shuttling between cytoplasm and nucleus in the PG is dependent on a functional circadian clock and a rhythmic PTTH input.

### Neuronal activity of PTTHn is required at the end of pupal development for the circadian gating of eclosion

To determine when during pupal development the rhythmic activity of PTTH is required for a circadian gating of emergence, we investigated the consequences on the timing of emergence of conditionally silencing the PTTHn. To do so, we expressed the inward rectifying K^+^ channel, Kir2.1, at different times of development using the temperature dependent TARGET system (McGuire et al., 2004). As shown in Fig. 5A and 5B, silencing the PTTHn during the embryonic and larval stages or during early pupal development did not affect the circadian rhythmicity of emergence. By contrast, silencing these neurons during the entire or only the second half of metamorphosis caused the expression of an arrhythmic pattern of emergence. These results indicate that the activity of PTTHn is required during the final stages of metamorphosis for the clock to impose circadian rhythmicity to the temporal pattern of adult emergence.

**Figure 5.**
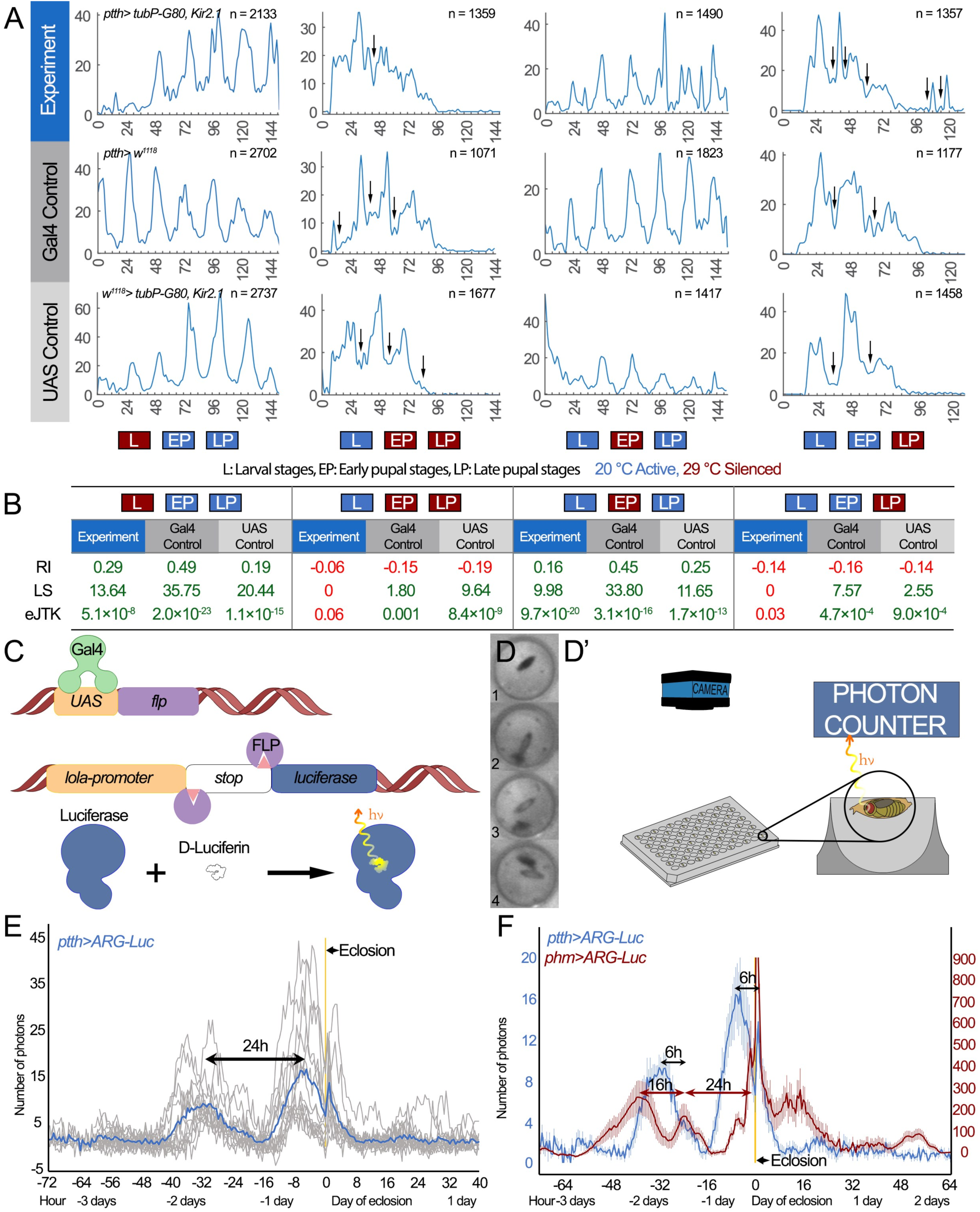
PTTHn activity is required during the end of pupal development. (A) Record of the time course of emergence in *ptth>tubP-Gal80^ts^, Kir2.1* flies with PTTHn conditionally silenced at 29°C during larval (L), entire pupal (EP+LP), early pupal (EP), and late pupal stages (LP) and in corresponding controls. Rhythmic records show peaks followed by valleys (arrows), which are separated by ∼24 hours. **(B)** Rhythmicity index (RI), Lomb-Scargle (LS) and eJTK_Cycle values of experiments shown in (A). Rhythmic records are coded in green, arrhythmic ones in red. Raising the temperature to 29°C during metamorphosis accelerates development, and when extended to the late pupal stage (LP) causes emergence to occur in only two to three peaks eclosion peaks. This renders autocorrelation inappropriate for rhythmicity analysis. Hence values are given for LS and eJTK_Cycle only. **(C-F)** *In vivo* imaging of PTTHn and PG activity prior to emergence. **(C-D’)** Schematic representation of the ARG-Luc system and *in vivo* imaging setup. (**D)** Images of intact animals captured at different stages: pharate [1]; eclosing [2]; eclosed; prior to [3], and after [4], wing expansion. **(E)** ARG-Luc activity of PTTHn for 14 individual flies plotted relative to the time of emergence (yellow line); blue line corresponds to the average trace. **(F)** Average ARG-Luc activity for the PTTHn (blue) and for the PG (red, n=22). Error bars indicate SEM. For PTTHn, 21 animals were recorded, but the records of only 14 were included in E and F; the remaining 7 animals were excluded because their signal did not change during the recording period, most likely because they were either located too far from the sensor or were not correctly oriented. The peaks immediately following eclosion in E and F are likely artifacts due to higher photon yields once flies have left their absorbing puparium.

In order to directly assess the activity of PTTHn and of the PG during the second half of metamorphosis, and considering that we are not able to perform *in vivo* Ca^2+^ imaging of the PG at this time, we turned to an Activity-Regulated Gene-Luciferase reporter (ARG-Luc)(Chen et al., 2016) for long-term monitoring of neuronal activity in intact developing pupae (Figure 5C and 5D). Strikingly, we found that PTTHn under DD conditions expresses a monophasic circadian rhythm of activity that started 2 days prior to adult emergence, with a second peak of activity preceding eclosion by 6h (Figure 5E and 5F). This timing is consistent with the peak in intracellular Ca^2+^ signaling detected in the middle of the subjective night in DD at the WPP stage (Fig. 2E), as well as with the increased signal detected just before emergence using the calcium-dependent sensor, CaLexA (Masuyama et al., 2012) (Fig. S6). In parallel experiments, we also used ARG-Luc to monitor the activity of the PG during pupal development. As shown in Fig. 5F, the PG expressed a peak of activity at around -25h and a large peak at around the time of eclosion, each lagging PTTHn activity by around 6h. In addition, our records showed an earlier peak of activity in the PG at around -40h that was not preceded by a peak in PTTHn activity, suggesting that activity in the PG depends on other factors in addition to PTTH but that its phase close to the time of emergence is set by PTTHn activity. (In the PG, increases in ARG-LUC signal may be due to changes in Ca^2+^ levels (Morioka et al., 2012) but we cannot rule out that they may be caused by a different mechanism; Chen et al., 2016.)

### PTTH-independent RTK signaling pathways contribute to the rhythm of the adult emergence

Although PTTH is the best known ecdysteroidogenic factor, the peak of activity detected in the PG 40h before eclosion preceded the first peak of activity in PTTHn, which may occur in response to other signals that act on the PG (Colombani et al., 2005; Cruz et al., 2020; Pan and O’Connor, 2021). In particular, a recent study reported that anaplastic lymphoma kinase (*Alk*) and PDGF and VEGF receptor-related (*Pvr*), two receptor tyrosine kinases (RTK) that are expressed in the PG, act in an additive manner to regulate the timing of metamorphosis (Pan and O’Connor, 2021). To determine whether these RTKs contribute to the rhythm of adult emergence, we determined the consequences on the timing of eclosion of knocking down or of expressing dominant-negative versions of these receptors, in the PG. As shown in Figures 6A-B and 6I, overexpression of a dominant-negative version of *Pvr* in the PG weakened the rhythm of the emergence relative to its control. A similar phenotype was produced by expressing two different *Alk* RNAi constructs (Figure 6C), as well as when both receptors were knocked down together (Figures 6D and 6I). Flies bearing only UAS-RNAi transgenes for *Alk* and *Pvr* expressed normal circadian rhythmicity of emergence (Figure S7A and S7B). It has been shown that Alk acts via both PI3K and Ras/Erk signaling pathways (Pan and O’Connor, 2021). Although we previously showed that Ras/Erk is critical for the rhythm of emergence (Selcho et al., 2017), we found that knockdown of *pi3k* in the PG did not affect the rhythm or period of emergence (Table S1). These results suggest that Alk contributes to the circadian control of eclosion exclusively via the Ras/Erk signaling pathway. Nevertheless, the contribution to the ERK pathway from receptors other than TORSO is comparatively minor since SPARK rhythmicity was not detected in the PG of PTTH null mutant animals (Fig. 3I) nor when *torso* was knocked down in the PG (Fig. 4I).

**Figure 6.**
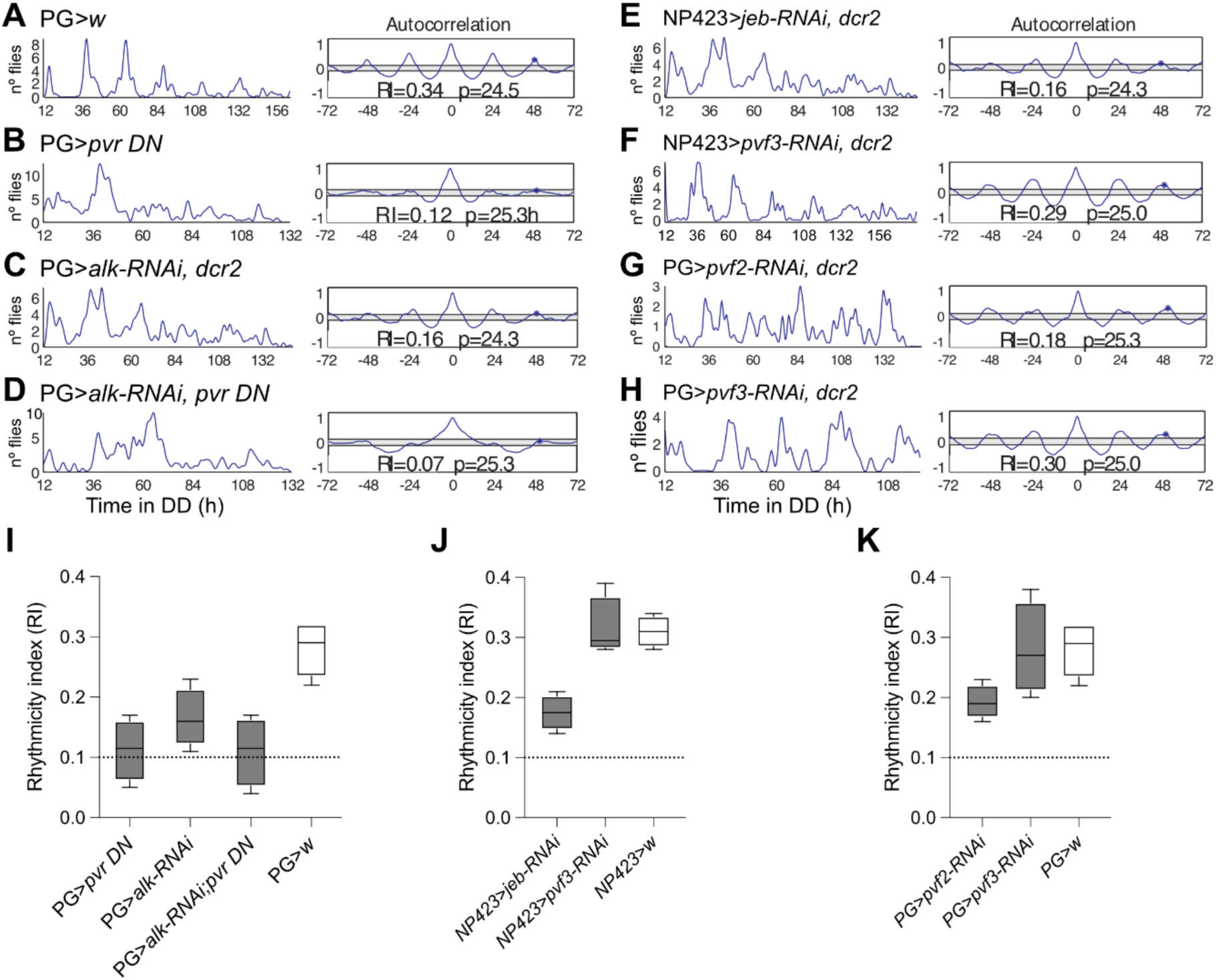
Alk and Pvr contribute to the circadian rhythmicity of adult emergence. **(A-H)** Records showing the time course of emergence under DD (left) and corresponding autocorrelation analysis (right) of a single population of flies expressing: in the PG, a dominant negative form of *Pvr* **(B)**, *Alk* RNAi **(C)**, simultaneous knockdown of PVR and ALK receptors **(D)**, *Pvf2* RNAi **(G)** and *Pvf3* RNAi **(H)**; and expressing in PTTHn: *jeb* RNAi **(E)** and *Pvf3* RNAi **(F)**. Periodicity (p, in hours) and associated rhythmicity index (RI) are indicated. **(I-K)** Average RI values for results shown in (A-H) and their respective controls; dashed line indicates cutoff below which records are considered arrhythmic. Different letters indicate statistically significant differences (*p < 0.05; one-way ANOVA, Tukey *post hoc* analysis.

Recent work has revealed that PTTHn express the Alk ligand, Jeb (Jelly belly), and one of three known ligands for Pvr, Pvf3, (Pan and O’Connor, 2021). Consistent with the consequences of knocking down the Alk receptor in the PG, knockdown of its ligand, Jeb, in PTTHn using the strong *NP423-GAL4* driver (Yamanaka et al., 2013) weakened the rhythmicity of emergence (Figures 6E and 6J). By contrast, the corresponding knockdown of *Pvf3* was without effect (Figure 6F and 6J). Finally, given that Pvf2 (another ligand for Pvr) and Pvf3 are expressed in the PG (Pan and O’Connor, 2021), we explored the possibility that these ligands might regulate the circadian rhythmicity of emergence in an autocrine manner. Interestingly, knockdown of *Pvf2* (but not of *Pvf3*) in the PG weakened the rhythmicity of adult eclosion (Figures 6G, 6H, and 6K) suggesting that *Pvf2* contributes to the circadian rhythm of emergence *via* an autocrine pathway. Together, these results reveal that PTTH/Torso axis is a major, but not exclusive, regulator of the rhythmicity of emergence. Indeed, our results indicate that Jeb/Alk and autocrine signaling involving Pvf2/Pvr also contribute to the rhythmicity of eclosion.

We also explored whether other ecdysteroidogenic signals including Allatostatin A (Deveci et al., 2019), Corazonin (Imura et al., 2020), insulin signaling (Colombani et al., 2005), as well as Gq proteins encoded by CG30054 and CG17760 (Yamanaka et al., 2015), might contribute to the circadian control of adult emergence. As shown Table S1, RNAi knockdown or null mutations of components of these signaling pathways in the PG did not affect the rhythmicity or the periodicity of adult eclosion.

Since peptidergic neurons often co-express small molecule neurotransmitters(Nassel, 2018), we also investigated whether any of such signaling molecules are involved in the circadian control of emergence. As shown in Suppl. Fig. S8, we found that PTTHn do not express the signature markers of signaling by glutamate (VGLUT), GABA (VGAT) or acetylcholine (ChAT), suggesting that PTTHn may be exclusively peptidergic. Finally, we found that stopping the clock located in the fat body(Xu et al., 2008) by overexpressing the dominant negative form of *cycle* gene (cyc[λ-.901]) (Tanoue et al., 2004) did not affect the timing of adult fly emergence (Table S1), indicating that the fat body, which also houses a circadian clock and could influence the PG via endocrine signaling, does not contribute to the circadian regulation of adult emergence.

## Discussion

Circadian clocks impose daily periodicities to behavior, physiology, and metabolism (Roenneberg and Merrow, 2005; Herzog, 2007; Pilorz et al., 2018). This control is mediated by the activity of a central clock and of peripheral clocks, which are housed in a variety of relevant organs. Although these various clocks are known to be synchronized to provide the organism with a unified time, how this coordination occurs is not fully understood. Here, we characterized the cellular and molecular mechanisms involved in coupling two circadian clocks in *Drosophila*, the central clock of the brain and the peripheral clock located in the PG, which together restrict the time of emergence of the adult fly to the early part of the day. We showed that time information is propagated from the central sLNv pacemaker neurons to PTTHn through a peptidergic, non-synaptic, connection mediated by sNPF, and is then translated into a circadian regulation of Ca^+2^ levels and PTTH accumulation in PTTHn. Although these features were measured in animals initiating metamorphosis (white puparium stage, WPP), we found that Ca^+2^ levels in PTTHn also varied in the pharate adult and peaked just before adult emergence. (The fact that PTTHn showed a similar timecourse in Ca^+2^ levels at both developmental stages also suggests that the WPP is a relevant stage for investigating the clock control of behaviors that occur many days later, at adult emergence). Importantly, we found that neuronal activity of PTTHn at the end of pupal development is necessary for the circadian gating of eclosion. The PTTHn then transmit time information to the PG, where we show that PTTH imposes a daily rhythm to the transcript levels of *torso* and ERK phosphorylation, a key component of PTTH transduction. Although PTTH plays a critical role in the coupling between the central and the PG clock, we demonstrate here that Jeb/Alk and Pvf2/Pvr signaling also contribute to regulate the rhythm of emergence, revealing that this circadian behavior is the result of timed signals from several RTKs that converge onto the PG.

### The transduction of the time signal in the PG is unexpectedly complex

PTTH accumulation in terminals and somatic Ca^2+^ rhythms have similar phases, suggesting that the increases in Ca^2+^ may cause PTTH release onto the PG. (However, the timing of PTTH release remains hypothetical and will need to be directly demonstrated). Beyond that point, our findings reveal a complex relationship between the timing of PTTH abundance in the terminals of PTTHn, and the transduction of PTTH action in the PG. Indeed, PTTH immunoreactivity in PTTHn terminals and somatic Ca^2+^ levels were lowest during the subjective day, coincident with the highest *torso* mRNA expression and a transient maximum of phosphoERK in the PG. In turn, PTTH levels were highest in the terminals of PTTHn during the middle of the night, a time when *torso* mRNA expression and phosphoERK levels in the PG reached their lowest levels. Assuming that TORSO receptor expression follows *torso* mRNA levels (there are currently no reagents to reliably measure TORSO protein levels *in situ*), this temporal relationship is consistent with an anti-cooperative model proposed for the PTTH/Torso interaction in the silkworm (Jenni et al., 2015), where the actions of the ligand are dampened by low receptors levels, leading to a low sustained output, whereas high receptor levels induce a transient output burst. A similar regulation has also been described in *Drosophila* and in mammalian cells (Puig and Tjian, 2005) for the insulin receptor, InR, another RTK family member. Indeed, under high nutrient conditions, a known inductor of insulin secretion, cells down-regulate *InR* expression via the transcription factor FOXO. A similar mechanism could also apply to the clock control of glucocorticoid (GC) production by the mammalian adrenal gland, where the clock of the adrenal gland regulates the sensitivity to adrenocorticotrophic hormone (ACTH) (Oster et al., 2006).

### Convergence of RTK signaling onto the PG

Across insects including *Drosophila* (Mele and Johnson, 2019), RTK signaling plays a major role in the control of growth and the timing of the molts. In the PG, PTTH-TORSO, Jeb-ALK, and Pvf/Pvr signaling, act to regulate the timing of ecdysteroid production (Pan and O’Connor, 2021), which causes the animal’s developmental transitions. Here we showed that all three RTK signaling pathways also contribute to the circadian control of emergence. Eclosion requires ecdysteroid titers to fall below a certain threshold level (Sláma, 1980; Schwartz and Truman, 1983; Zitnan and Adams, 2012). However, the timing of emergence is not determined by the time when ecdysteroid titers drop below this threshold (Handler, 1982; Lavrynenko et al., 2015). Instead, it is PTTH (this work) and ecdysteroid signaling (Mark et al., 2021) that control the circadian gating of eclosion in *Drosophila* by committing the insect to complete metamorphosis around 14h before emergence (Mark et al., 2021). Since we observed peaks of activity in PTTHn around 31h and 6h before emergence, it may be the timing of the first rather than the second peak of activity in PTTHn that is critical for determining the timing of emergence. In this scenario, PTTH/Jeb signaling may not only promote ecdysteroid production, but also regulate the phase of the PG clock. Although our study does not provide mechanistic evidence for this scenario, it has been shown that RTK/MAPK signaling serves as both an output and an input pathway to the mammalian circadian clock (Goldsmith and Bell-Pedersen, 2013). Furthermore, a mutation in *Alk* was found to significantly alter the circadian period of locomotor activity in *Drosophila* (Kumar et al., 2021). In addition, it is known that *Pvr* and *Alk* expression are under circadian control in ventral and dorsal lateral clock neurons in the brain of the adult fly (Abruzzi et al., 2017), suggesting that these RTKs are downstream of the clock. It is therefore possible that the central-clock driven PTTHn-derived RTK signaling ensures proper eclosion timing by aligning the phases of the central and PG clocks.

Insulins are another class of ligands that act on the PG via an RTK (Colombani et al., 2005). An RNAseq study reported that insulin signaling modulates the expression of clock genes in the PG, which in turn regulates *torso* expression (Di Cara and King-Jones, 2016). Nevertheless, we found that insulin signaling in the PG does not affect the rhythm of emergence. Thus, not all signaling pathways mediated by RTKs are involved in the circadian control of adult emergence. Similarly, even though ALK acts through Ras/ERK and PI3K signaling pathways in the PG (Pan and O’Connor, 2021), we showed that PI3K signaling is not involved in the circadian gating of emergence, suggesting that, similar to TORSO, ALK and PVR contribute to the rhythm of emergence via the Ras/ERK pathway. Interestingly, the involvement of multiple signals in the transmission of the time information is consistent with the general principle of coherent coupling between multiple oscillators, which helps to stabilize and strengthen period, phase distribution, and amplitude in the SCN, and between central and peripheral body clocks (Pilorz et al., 2018; Schmal et al., 2018).

### Autonomous role of the PG clock in the timing of eclosion

We found that knockdown of the ligand, Pvf2, in the PG itself reduced the rhythmicity of emergence. This identifies elements of the circadian control of emergence that are autonomous to the PG, and may be responsible for the peak of ARC-LUC activity detected in the PG two days prior to eclosion, which occurred before the activation of PTTHn. These findings may explain why functional clocks in both the central brain and the PG are required for the daily gating of eclosion (Myers et al., 2003). Indeed, since *torso* levels do not cycle in the absence of PTTH, it appears that the PG clock does not control the sensitivity to PTTH, raising the question of what role the clock of the PG plays in regulating the timing of adult emergence. In addition, it is intriguing that the peak of activity in the PG occurs at eclosion because it is undergoing apoptosis at this time (Dai and Gilbert, 1991), raising the possibility that apoptosis is part of a sequence of physiological events that are regulated by the circadian clock and are critical for the timing of eclosion Much is currently known about how the circadian clock functions in a variety of multicellular organisms (Bell-Pedersen et al., 2005). By contrast, much less is known about how the clocks housed in different organs are coordinated. Interestingly, although animal clocks are universally controlled by intracellular transcriptional/translational feedback loops (Hardin, 2011; Takahashi, 2017), the mechanisms that mediate the coupling of clocks appears to be diverse and fine grained, such that in some cases some genes within a peripheral clock cycle autonomously whereas the cycling of others depends on a central input (Xu et al., 2011; Erion et al., 2016; Versteven et al., 2020). In addition, the relationship between central and peripheral clocks is even more complex if the integration of external stimuli such as light, temperature or feeding is considered (Ivanchenko et al., 2001; Glaser and Stanewsky, 2005; Mohawk et al., 2012; Ito and Tomioka, 2016). In the case of the circadian gating of adult emergence, we provide here a detailed analysis of the mechanism that couples a central and a peripheral clock, which may provide principles that apply to other circadian clocks. Knowing how clocks are coordinated will be critical to understand how circadian rhythmicity is generated at the cell, systems, and organism levels.

## Acknowledgments

This work was funded by ANID Graduate Fellowship #21180133 (to JC), the Deutsche Forschungsgemeinschaft (DFG WE 2652/7-1, to CW and STA 421/8-1 to RS), and FONDECYT (National Fund for Scientific and Technological Development, Chile) grant #1221270 (to JE); and the ANID – Millennium Science Initiative Program “Centro Interdisciplinario de Neurociencia de Valparaiso (CINV)” grant P09-022-F (to JE). We thank Sebastian Cachero, Alisson Gontijo, Sebastian Hückesfeld and Michael Pankratz, Michael O’Connor, Tom Kornberg, Matthias Schlichting and Michael Rosbash, Peter Soba, Paul Taghert and Meet Zandawala, for the kind gift of flies and antisera. We thank Alexander Veh for help with eclosion assays.

## EXPERIMENTAL MODEL AND SUBJECT DETAILS

### Fly stocks and husbandry

Flies were raised on standard cornmeal/yeast media and maintained at room temperature (20 to 22 °C) under a 12h:12h LD schedule. All UAS and GAL4 stocks have previously been described; unless noted, they were obtained from the Bloomington *Drosophila* stock center (Bloomington, Indiana, USA; BL) or the Vienna *Drosophila* Resource Center (Vienna, Austria; VDRC). They included: wildtype (Canton-S strain), *white*[1118], *ptth*-GAL4(45A3,117b3)(McBrayer et al., 2007), *np423-*GAL4(Yamanaka et al., 2013), *phm*-GAL4 (obtained from Michael O’Connor), *Pdf*-GAL4 (obtained from Paul Taghert), *lsp*-GAL4 (BL6357), UAS-cyc[Δ901](Tanoue et al., 2004), 20xUAS-IVS-GCaMP6m(Chen et al., 2013), UAS-ANF-GFP(Husain and Ewer, 2004), UAS-EGFR (BL9535), UAS-*Pvr* DN (BL58431), *ptth delta*(Shimell et al., 2018), UAS-EGFP-Kir2.1(Baines et al., 2001) (obtained from Sebastian Hückesfeld and Michael Pankratz), tubP-Gal80^ts^ (BL7019), lexAop-nSyb-spGFP1-10, UAS-CD4-spGFP11(Macpherson et al., 2015) (obtained from Peter Soba), LexAop2-Syb.GFP.P10 LexAop2-Syb.GFP^N146I^.TEV^T173V^ LexAop2-QF2.V5.hSNAP25.HIVNES.Syx1A/CyO, 20XUAS-B3R.PEST UAS(B3RT.B2)BoNTA QUAS-mtdTomato-3xHA/TM6B, Tb^1^ (obtained from Sebastian Cachero)(Cachero et al., 2020), CaLexA (BL66542), 20XUAS-flp; lola-frt-stop-frt-luc(Chen et al., 2016) (obtained from Matthias Schlichting and Michael Rosbash), hsFLP UAS-mCD8::GFP.L QUAS-mtdTomato-3xHA; trans-TangoMkII/SM6b (BL95317), QUAS-nlsDsRed (obtained from Meet Zandawala)(Snell et al., 2022), *dilp2, 3, 5* and *dilp 7* mutants(Gronke et al., 2010) (BL30889 and kind gift of Peter Soba), *dilp8* and *Lgr3* mutants(Garelli et al., 2012; Garelli et al., 2015) (kind gift of Alisson Gontijo). Flies bearing UAS-SPARK (Zhang et al., 2018) and UAS-SPARK (TèA)(Zhang et al., 2018) were provided Tom Kornberg, All genotypes involving the use of UAS-RNAi transgenes included a copy of a UAS-*dcr2* transgene. When performing RNAi knockdown, several UAS-RNAi lines were tested and the one producing the most severe phenotype was used. The lines used and their source is indicated below; the line chosen for the results reported here are indicated in bold: *IP3R* (Inositol 1,4,5, -tris-phosphate receptor; CG1063): **BL#25937**, VDRC#6486; *Dmca1A* (*cacophony* or Calcium-channel protein α1 subunit A; CG1522): **BL#27244**, VDRC #104168; *Dmca1D* (*Calcium-channel protein α1 subunit D*; CG4894): BL#25830; *SERCA* (*sarcoendoplasmic reticulum calcium ATPase* or *Calcium ATPase 60A*; CG3725): **VDRC#4474**, VDRC#107446, BL#25928; *RyR* (*ryanodine receptor*): **VDRC#109631**, BL#29445, *CASK* (*calmodulin-dependent protein kinase activity*; CG6703): **BL#27556**, BL#35309; *PKC* (*protein kinase C*); **VDRC#27699**, BL27491; *AstAR1*: **VDRC#39222**, VDRC#48495; *CrzR*: **VDRC108506**, VDRC#44310; *torso*: VDRC#36280; *Alk*: **BL#27518**, VDRC#107083; *Pvf2:* **BL#61195**, VDRC#7629 *Pvf3*: VDRC#373933; *Jeb:* **VDRC**#**103047**, CG30054: **BL#51823**, VDRC#4643; CG17760: **BL#64919**, VDRC#52308; *InR*: VDRC#992; PI3K: VDRC#107390; PTEN: VDRC#35731.

## METHOD DETAILS

### Emergence monitoring

Flies were raised at 20°C under 12 h light and 12 h darkness (LD 12:12). After 12–17 days, pupae were collected and fixed on eclosion plates with Elmer’s glue or methyl cellulose glue (Tapetenkleister #389, Auro, Braunschweig, Germany) and mounted on Trikinetics eclosion monitors (Trikinetics, MA, USA). Emergence was then monitored for 7 days at 20°C under DD conditions, in climate-and light-controlled chambers. Rhythmicity of eclosion profiles was analyzed using MATLAB (MathWorks, Inc., Natick, USA) and the appropriate Matlab toolbox(Levine et al., 2002). Using autocorrelation analyses, records were considered rhythmic if the rhythmicity index (RI) was ≥0.3, weakly rhythmic if 0.1:sRI<0.3 and arrhythmic if RI<0.1 (Sundram et al., 2012). In the case of experiments in which the temperature was increased to 29°C during metamorphosis, the increase in temperature causes a faster development, leading to a reduced number of eclosion peaks. In such cases autocorrelation analysis produced low RI values for controls so Lomb-Scargle (LS) (Ruf, 1999) and eJTK (Hughes et al., 2010) analyses were used instead.

### Ex vivo measurements of Ca^+2^ and phosphorylated ERK signals

Animals were reared under 12h:12h LD conditions with lights-on at noon or midnight. Animals used were always at the start of metamorphosis (white pre-pupal stage, WPP). Thus, developmental stage was kept constant; the only variable was time of day. For measurements made under DD conditions, cultures were wrapped in aluminum foil at lights-off and maintained covered until the desired time. In order to determine Ca^+2^ levels in PTTHn, we measured the GFP fluorescence of these neurons expressing the Ca^+2^ sensor, GCaMP6m. For this, WPP stage animals were collected, and their brains dissected under ice-cold calcium-free fly saline (46mM NaCl, 5mM KCl, and 10mM Tris pH 7.2). They were then mounted on slides coated with poly-lysine in an Attofluor chamber (A-7816, Invitrogen) and imaged on an Olympus Spinning Disc microscope using an UMPlanFI 20X/0.50 water immersion objective and CellSens software. Samples were visualized using the GFP filter (excitation 485nm and emission at 515nm) and imaged using a Hamamatsu camera (model ORCA IR2). Regions of interest (ROIs) were drawn over the cell bodies of PTTHn. Background was subtracted for every ROI and the changes in fluorescence intensity were calculated as:

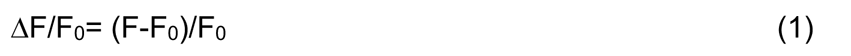

Where F is the fluorescence in the cell bodies of PTTHn and F_0_ is the baseline fluorescence value in the brain. Mean fluorescence intensities were measured using Fiji (Schindelin et al., 2012). Data were analyzed in Microsoft Excel.

Levels of phosphorylated ERK in the PG were determined by driving expression of the SPARK sensor in the PG and measuring the GFP fluorescence emitted by individual PG cells. PGs dissections and imaging were performed as described above. For quantitative analysis of the SPARK signal, images were processed using Fiji(Schindelin et al., 2012). The droplets included in the analyses were selected using the ‘‘threshold” function, considering only those with a size larger than that of the residual droplets observed when the mutant (inactive) version of ERK-SPARK was expressed in the PG. Normalized SPARK was calculated as:

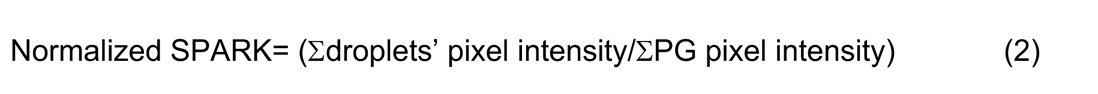

The sum of droplet pixel fluorescence intensity and the PG pixel intensity were calculated using Analyze Particle function in the software. For the quantification of nuclear droplets, only droplets with a clear nuclear localization were considered; droplets in a region between nucleus/cytoplasm were excluded.

### CaLexA Ca^2+^ imaging

Eggs of *ptth>CaLexA* flies were collected for five hours and kept at 12:12 LD, 25°C, and 65% humidity. Pupal brains were dissected at pupal stage P14 in 4-hour intervals at ZT0, 4, 8, 12, 16, and 20, and immediately fixed and stained for GFP-and PTTH-immunoreactivity as outlined above. After mounting, the brains were imaged under a confocal microscope using the same settings for all preparations. Staining intensities were determined using Fiji.

### ARG-Luc imaging

*ptth>*20X*UAS-flp; lola-frt-stop-frt-luc* larvae were raised in the dark on normal food supplemented with 15 mM luciferin (Carbosynth Ltd., Newbury, UK) until pupariation. A 1h light pulse was given at ZT0 to synchronize the clocks of the animals used. The next day puparia were transferred to light during the subjective photophase and washed from the bottles, dried, and glued on adhesive transparent plastic sheets (TopSeal™ A Plus (PerkinElmer LAS, Rodgau, Germany) on top of a 96 well OptiPlate™ (PerkinElmer LAS, Rodgau, Germany). Three days before the estimated time of eclosion, puparia were glued on every other well to minimize light spill-over. Bioluminescence was recorded every 30 min in DD at 25°C and 65% humidity using a TopCount Multiplate Reader (Perkin Elmer). An sCMOS camera (DMK33UX178, ImagingSource, Bremen, Germany) on top of the plate recorded a picture every 5 minutes under red light to monitor the eclosion state. Rhythmicity of the ARG-Luc signals was analyzed using MESA and JTK_Cycle implemented in BioDare2 (Zielinski et al., 2014).

### PTTH antiserum production

Rabbit polyclonal PTTH antiserum to PTTH was produced against the synthetic peptide CQSDHPYSWMNKDQPWamide (residues 207-221, with an N-terminal Cys added), not amidated which was coupled to thyroglobulin via the N-terminal Cys residue using maleimide. The immunization was carried out by Pineda-Antikörper Service (Berlin, Germany) and lasted over 3 months, with a booster before the final bleed.

### Immunocytochemistry

Dissected brains or PGs were fixed in 4% paraformaldehyde for 15 minutes at room temperature (RT). Tissues were washed in PBS containing 0.3% TritonX-100 (PBT), blocked with 5% normal goat serum (NGS) in PBT, and incubated overnight at 4°C in primary antibodies. Primary antibodies used are listed in Table 1. Tissues were then washed in PBT and incubated for 2h at RT in secondary antibody, rinsed three times in PBS, and mounted on a poly-L-lysine coated coverslip using Fluoromont-G T (Electron Microscopy Science, USA). Alexa Fluor conjugated secondary antibodies (Alexa Fluor 488 conjugated goat anti-guinea pig, and goat anti-rabbit were from Invitrogen, MA, USA) were used at 1:500.

Tissues were imaged on an Olympus Spinning Disc microscope using either UPLFLN20X/0.5NA or UPLSAP040X/0.9NA objectives, and were analyzed using ImageJ. Mean fluorescence intensity in sLNv terminals or PTTH signal in the PTTHn cell bodies was quantified and the signal from brain areas devoid of specific immunoreactivity was subtracted as background. For the quantification of the number of PTTH immunoreactive boutons on the PG, boutons were selected using the “threshold” function and quantified using ImageJ’s “analyze particles” function.

### Immunocytochemistry for (chemo)connectomics, BacTrace, and syb-GRASP

Brains of P14 pupal stage adults were dissected in HL3.1 fly ringer (70 mM NaCl, 5mM KCl, 1.5mM CaCl_2_.2H_2_O, 4mM MgCl_2_.6H_2_O, 10mM NaHCO_3_, 115mM sucrose, 5mM Trehalose.2H_2_O, and 5mM HEPES(Feng et al., 2004)) and collected on ice. The HL3.1 solution was then replaced by 4% paraformaldehyde in phosphate-buffered saline (100 mM PBS, pH 7.4) and fixed for 30 minutes. Tissues were then washed three times with PBS with 0.3% Triton-X100 (PBT) for 10 minutes, followed by blocking in 5% normal goat serum (NGS) in PBT for 90 minutes. After blocking, the tissues were transferred into primary antibody diluted in PBT containing 3% NGS for 3 days at 4°C on a shaker, washed 6×20 min with PBT, and incubated for 24h at 4°C with secondary antibody in PBT containing 3% NGS. Finally, the tissues were washed 4×10min with PBT and 3×10min with PBS, and mounted on poly-lysine-coated slides (Thermo Scientific) with Vectashield (Vector Laboratories, California, United States), and stored at 4°C until imaged.

To visualize the syb-GRASP signal, freshly dissected brains were briefly (5-7s) depolarized three times using HL3.1 containing 70 mM KCl, then returned for 10 min in HL3.1 to allow reconstruction of GFP at active synaptic sites prior to fixation. For *trans-Tango* MkII, animals were raised at 18°C until P14 in order to maximize the accumulation of signal in post-synaptic neurons. In both cases, tissues were fixed for 30 minutes with 4% PFA in PBS and processed as described above.

Tissues were imaged on a Leica TCS SPE confocal laser scanning microscope (Leica Microsystems, Wetzlar, Germany) equipped with 20x and 40x high aperture immersion objectives.

### qRT-PCR

For each timepoint point, 40 PGs from WPP animals were dissected under ice-cold phosphate-buffered saline (PBS) and stored in RNAlater solution (Qiagen). Total RNA was extracted using Trizol reagent (Invitrogen, Carlsbad, CA, USA) following manufacturer’s instructions, and resuspended in 20 ml of RNAse-free water.

cDNA synthesis was carried out using SuperScript II Reverse Transcriptase (Invitrogen) following manufacturer’s instructions. One μg of RNA was first treated for 15 min at room temperature with 1 μl DNAse I (2U μl^-1^), 10X DNAse I Buffer (100 mM Tris, pH 7.5; 25 mM MgCl_2_; 5 mM CaCl_2_); DNAse I was then inactivated adding 1.5 μl 25 mM EDTA and the reaction incubated for 10 min at 65 °C. Oligo dT (0.5 μg) and dNTP’s (10mM each) were then added to each sample, incubated at 65 °C for 5min, then placed on ice. Four μl of 5X First-Strand Buffer (250 mM Tris-HCl, pH 8.3; 375 mM KCl; 15 mM MgCl2) and 2 μl of DTT (100 mM) were then added, and samples incubated for 2min at 42 °C. Finally, 1 μl of Reverse Transcriptase was added, and the reaction incubated at 42 °C for 50 min, followed by 15 min at 70 °C. Control reactions were processed in parallel except that the Reverse Transcriptase was omitted.

Quantitative PCR with reverse transcription (qRT-PCR) was carried out using *torso*-specific primers suitable for qRT-PCR; *rp49* was used as reference (see Supplementary Table 1 for the sequence of the primers used). Reaction mix included 5 μl Maxima SYBR Green/ROX qPCR Master Mix 2X (Thermo Fisher Scientific Inc., Waltham, MA, USA), 0.25 ml primers (10 μM), 1 μl cDNA, and 3.5 μl nuclease-free water. PCRs were carried out using an Agilent Mx3000P QPCR System thermocyler (Santa Clara, CA, USA) using the following regime: 25 °C for 5 s, 95 °C for 5 min, followed by 40 cycles of 15 s at 95 °C, 15 s at 60 °C and 15 s at 72 °C. A melting curve was done at the end of the reaction, which consisted of 10 s at 95 °C, 5 s at 25 °C, 5 s at 70 °C, and ending with 1 s at 95 °C. All analyses were carried out using MxPro QPCR Software (Agilent, Santa Clara, CA, USA). At least three independently isolated cDNAs were used, and each cDNA was qRT-PCR amplified in triplicate.

### Quantification and statistical analyses

Parametric datasets were first analyzed for normal distribution using the D’Agostino-Pearson or Shapiro-Wilk normality test using Prism 8.0 (GraphPad, USA), then analyzed using unpaired Student’s t-test or one-way ANOVA followed by Tukey’s multiple comparisons tests. For non-parametric datasets, we used Kruskal-Wallis tests followed by Dunn’s multiple comparisons tests. The results of all statistical tests are shown in Table S3.

## Material Availability

This study generated a new polyclonal rabbit antiserum against PTTH. Requests should be directed to D.R.N.

## Data and code availability

All relevant data are included in this report. No codes were generated in this study.

