## Supplemental files for "Timed receptor tyrosine kinase signaling couples the central and a peripheral circadian clock in *Drosophila*"

### SUPPLEMENTARY INFORMATION

#### SUPPLEMENTARY FIGURES

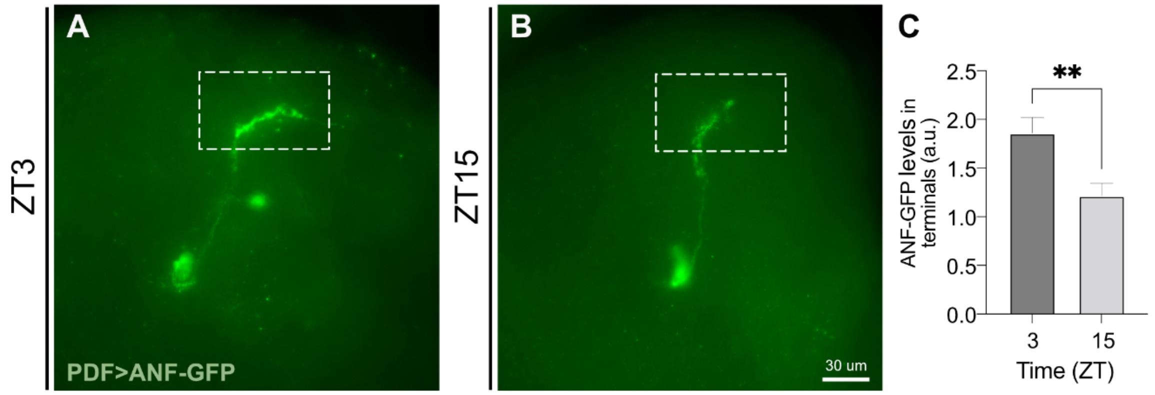


**Figure S1. Rhythm of neuropeptide accumulation in sLNv terminals. (A-B)** ANF-GFP signal in PDF neurons from wildtype animals at the WPP stage at ZT3 **(A)** and ZT15. **(B)**. Dashed box indicates the sLNv terminals. **(C)** Average ANF-GFP fluorescence levels in the sLNv terminals. Eight to 10 brains were analyzed per time-point (**: $p<0.05$; Student t-test).

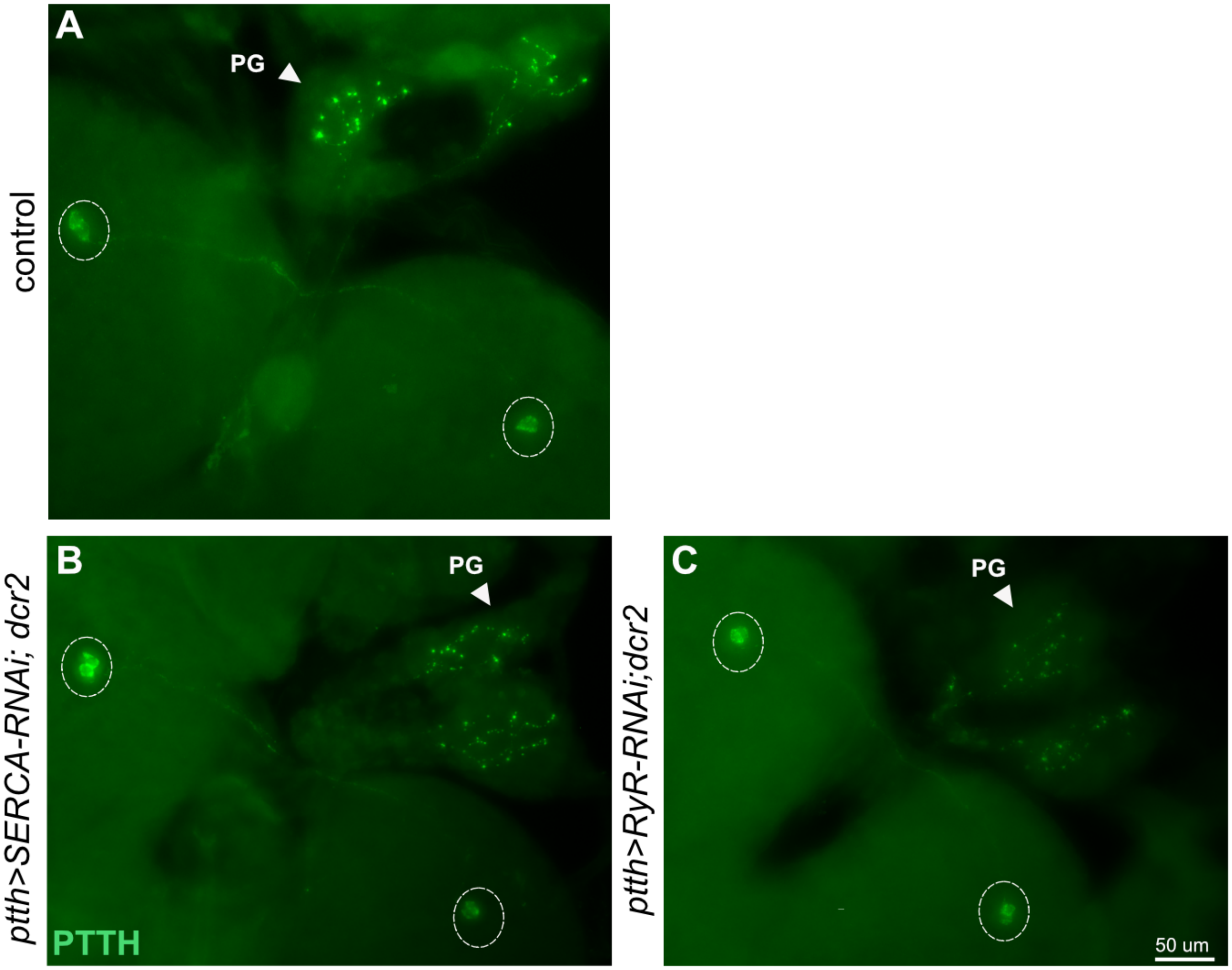


**Figure S2. Morphology of PTTHn is not grossly affected by knockdown of SERCA or RyR. (A-C)** Whole mount of brain-PG complex immunostained for PTTH from control (*ptth>w*) **(A)**, *ptth>SERCA-RNAi;dcr2* **(B)** and *ptth>RyR-RNAi,dcr2* **(B)** animals at the WPP stage. Dashed circles indicate PTTHn cell bodies. Bright spots on PG (arrowhead) correspond to the terminals of PTTHn. (Images were not all acquired with the same exposure time and thus, fluorescence levels are not directly comparable.)

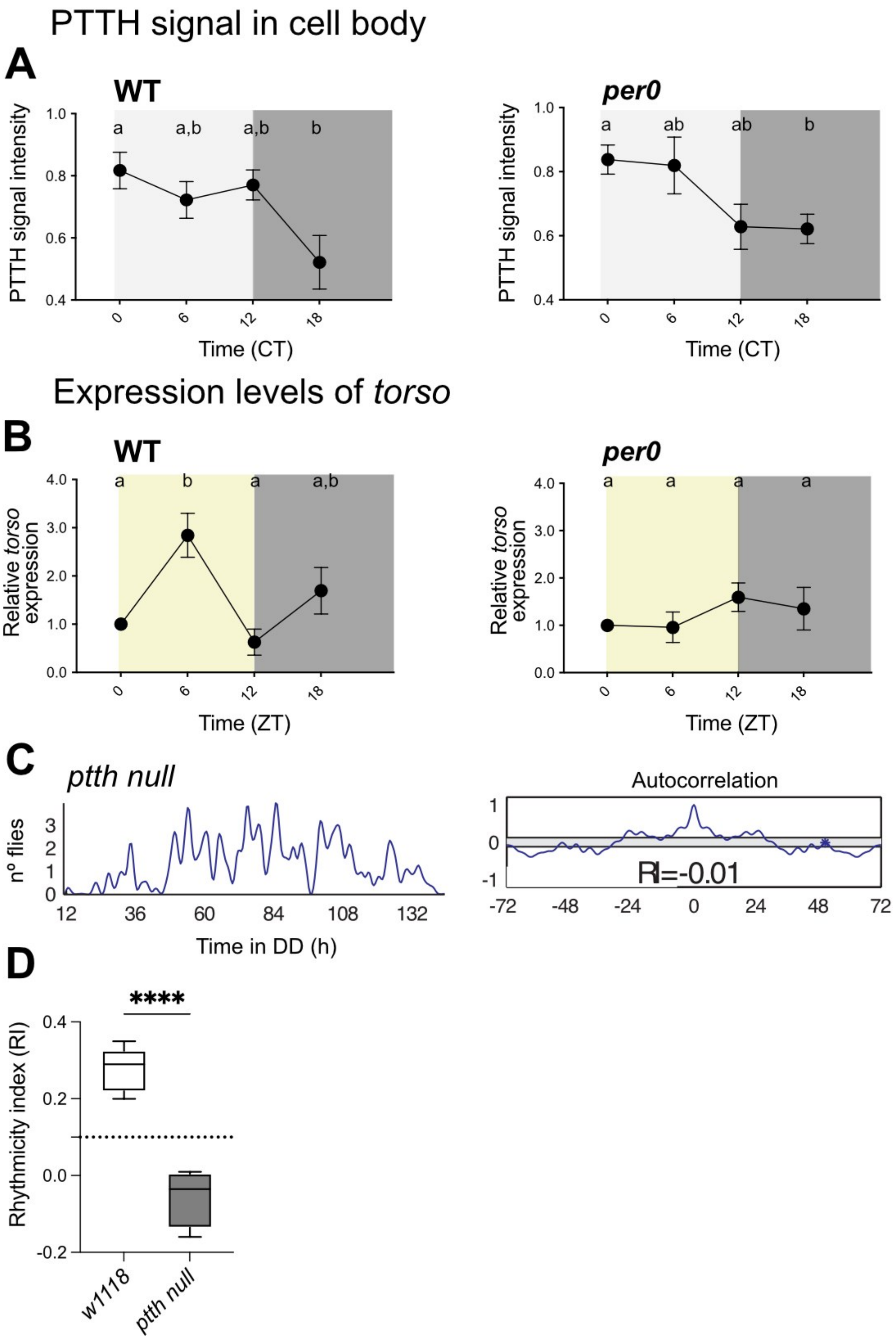
PTTH signal in cell body
A
WT
per0
PTTH signal intensity
a
a,b
a,b
b
ab
ab
b
1.0
0.8
0.6
0.4
0
6
12
18
Time (CT)
Expression levels of torso
B
WT
per0
Relative torso expression
4.0
3.0
2.0
1.0
0.0
Time (ZT)
C
ptth null
nº flies
Time in DD (h)
12
36
60
84
108
132
Autocorrelation
RI=-0.01
-72
-48
-24
0
24
48
72
D
****
Rhythmicity index (RI)
0.4
0.2
0.0
-0.2
w1118
ptth null

**Figure S3. Daily rhythm in PTTH levels in PTTH neurons and *torso* expression in the PG.** **(A)** Quantification of PTTH immunoreactivity in the cell bodies of PTTHn in wildtype (left) and *per[0]* mutant (right) animals at the WPP stage at different times of (subjective) day under DD conditions. **(B)** qRT-PCR analysis of *torso* expression in PGs dissected from WPP stage wildtype (left) and *per[0]* mutant (right) animals under LD conditions. *rp49* was used as reference gene. **(C)** Record showing timecourse of emergence of a single population of flies under DD conditions (left) and corresponding autocorrelation analysis (right) in *ptth* null mutant animals. Rhythmicity index (RI) associated is indicated. **(D)** Average values for RI (left) for *ptth null* mutant animals and controls (****: $p<0.0001$; Student t-test).

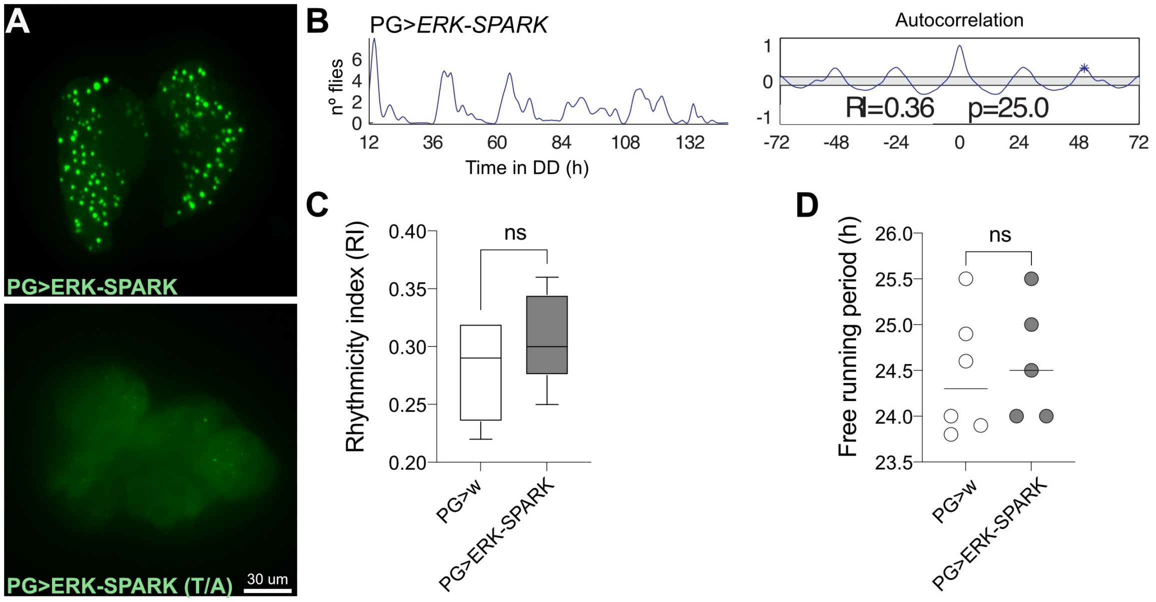


**Figure S4. SPARK reports ERK activity in the PG without affecting the circadian control of adult emergence.** **(A)** PG whole mount of animals expressing ERK-SPARK sensor (top) or ERK-unresponsive mutant sensor (bottom). **(B)** Record showing time course of emergence of a single population of flies under DD conditions (left) and corresponding autocorrelation analysis (right) of a population of flies expressing ERK-SPARK in PG. Periodicity (p, in hours) and associated rhythmicity index (RI) are indicated. **(C-D)** Average values for RI **(C)** and periodicity **(D)** for PG>*ERK-SPARK* and for control. Different points are results from a separate experiment (ns: $p>0.05$; Student t-test).

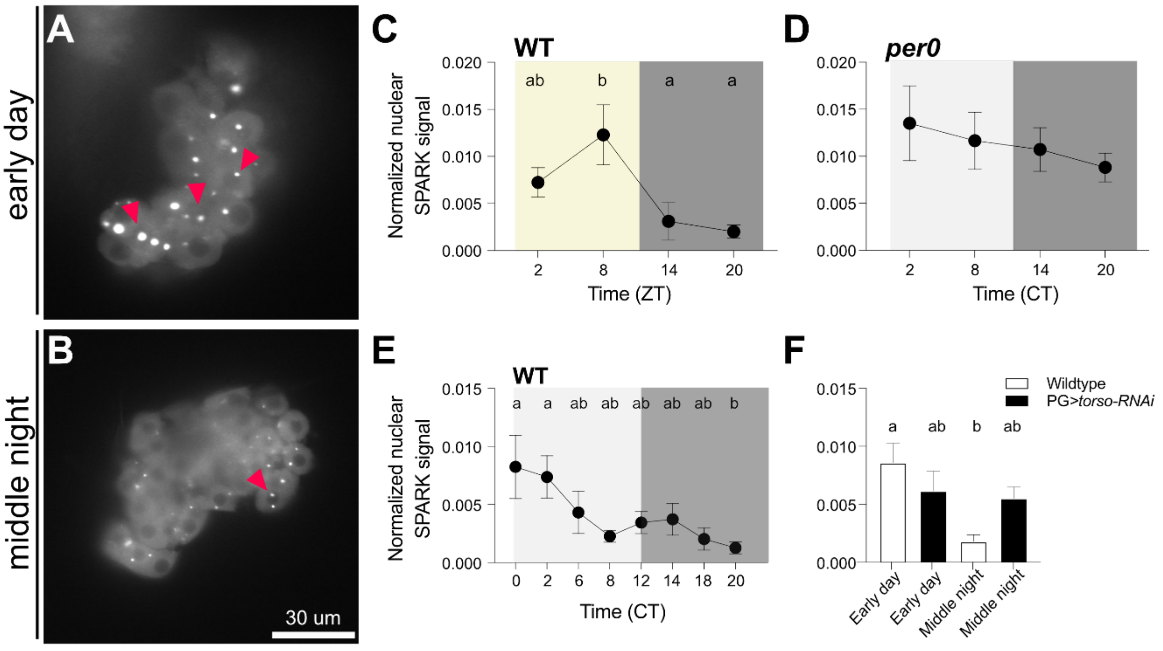


**Figure S5. Nuclear localization of phosphorylated ERK in the PG exhibits a daily rhythm.** **(A-B)** PG whole mount of animals expressing ERK-SPARK in the PG (*phm>ERK-SPARK*) at CT0-2 (early subjective day; **A**) and CT18-20 (middle of subjective night; **B**). Red arrowheads point to droplets with nuclear localization. **(C-E)** Average of normalized nuclear SPARK signal at different times of day in wildtype under LD **(C)** and DD **(E)** conditions, and in *per0* mutants under DD condition **(D)**. Different letters indicate statistically different groups ($p<0.05$; ANOVA, Dunn's multiple comparison analyses). **(F)** Average of normalized nuclear SPARK signal in wildtype and PG>*torso-RNAi* animals at CT0-2 (early subjective day) and CT18-20 (middle of subjective night). 8-10 brains were analyzed per time-point. Different letters indicate statistically different groups ($p<0.05$; one-way ANOVA, Tukey's post hoc multiple comparison analyses).

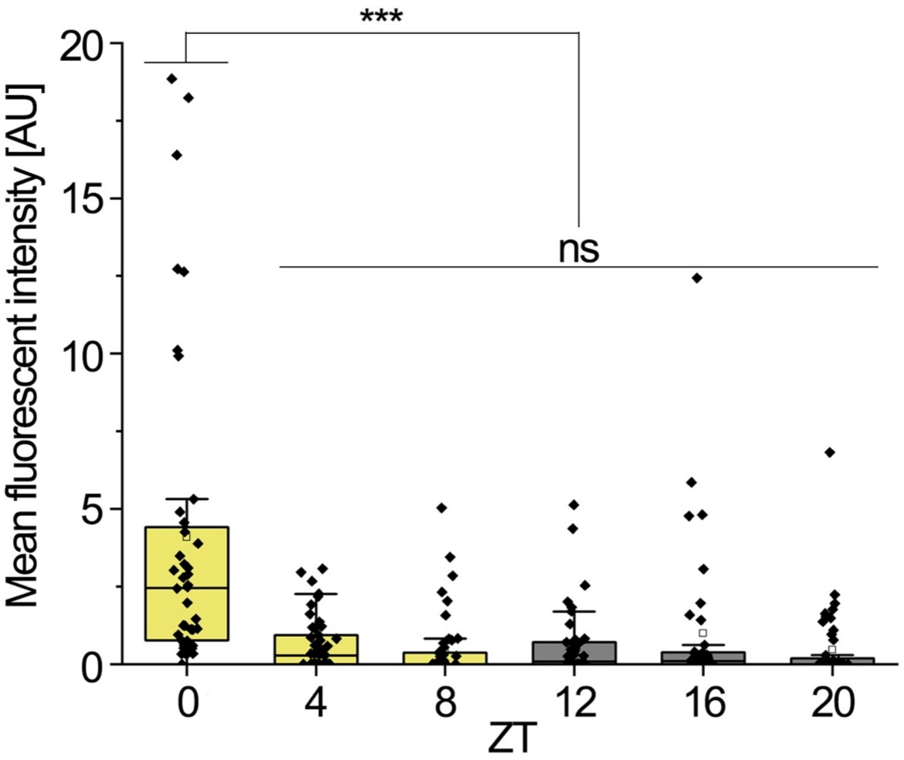


**Figure S6. $Ca^{2+}$-dependent CaLexA signals from the PTTHn suggests a peak of PTTHn activity preceding the emergence peak at dawn.** *ptth>CaLexA* flies were raised in 12:12 LD condition and brains from pharate adults were dissected in 4-hour intervals in the light phase (yellow) or under red light in the dark phase (gray). The highest GFP signal occurred at the expected time of emergence.

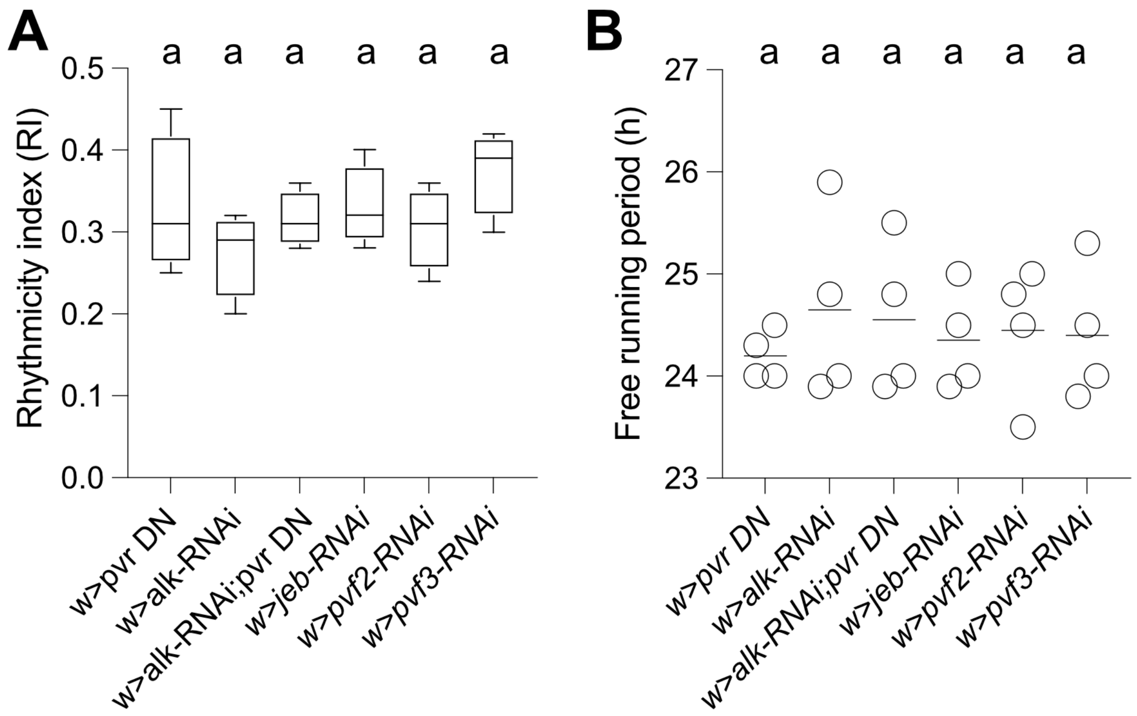


**Figure S7. Flies bearing single copies of UAS-RNAi insertions for *Alk* or *Pvr* signaling express normal circadian rhythmicity of eclosion. (A)** Average rhythmicity index (RI) values (± SEM) of flies heterozygous for UAS-RNAi transgenes for *Pvr*, *Alk*, *Alk*;*Pvr*, *jeb, pvf2*, and *pvf3*. Same letters indicate that there are no statistically significant differences between groups (one-way ANOVA, Tukey's post hoc multiple comparison analyses). **(B)** Free-running period (h) values for genotypes indicated in (A); each point represents the results from a separate experiment; horizontal lines indicate the average. Same letters indicate that there are no statistically significant differences between groups (one-way ANOVA, Tukey's post hoc multiple comparison analyses

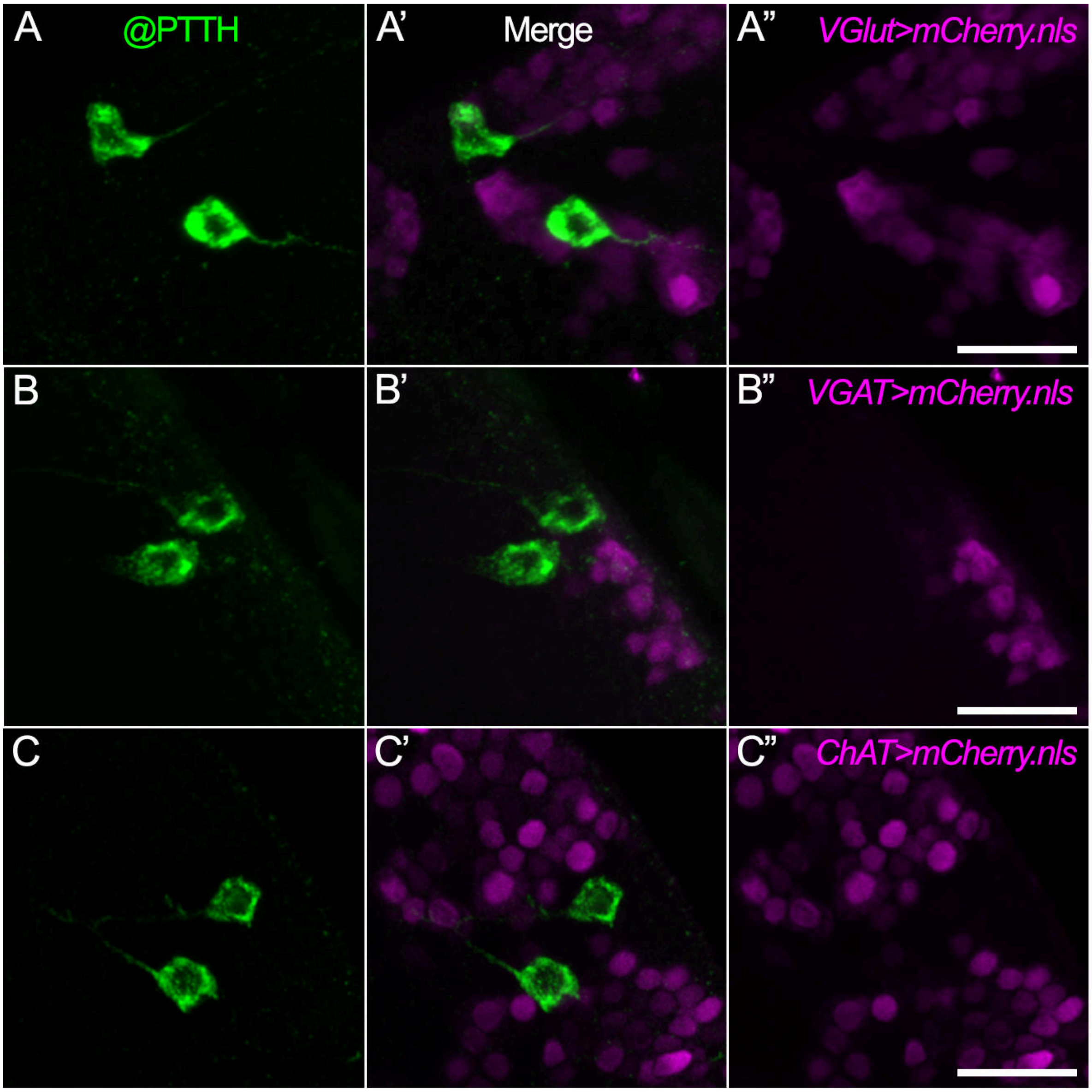


**Figure S8. PTTHn do not express the neurotransmitters glutamate, GABA or acetylcholine. (A")** Expression pattern of driver lines for vesicular glutamate transporter (*VGlut-Gal4*), **(B")** vesicular GABA Transporter (*VGAT-Gal4*) and **(C")** choline acetyltransferase (*ChAT-Gal4*). In all cases, there was no colocalization between the PTTHn (A, B, C) and labeling for glutamatergic (A'), GABAergic (B'), or cholinergic neurons (C'), suggesting that the PTTHn are exclusively peptidergic and do not signal via classic neurotransmitters. Scale bars*:* 20 *µm*.

### SUPPLEMENTARY TABLES

#### Table S1. Adult emergence rhythmicity phenotypes following genetic manipulations in the PTTH/PG axis or in the clock function.

| Genotype | Associated biological process | N (n) | RI ± SEM | Period (h) ± SEM |
|---|---|---|---|---|
| *ptth>GCaMP6M* | $Ca^{+2}$ imaging | 3 | 0.32 ± 0.02 | 24.1 ± 0.20 |
| *ptth>AstAR1*-RNAi | PTTH secretion (Deveci et al., 2019) | 5 | 0.30 ± 0.04 | 24.6 ± 0.22 |
| *ptth>CrzR*-RNAi | PTTHn activity (Imura et al., 2020) | 5 | 0.35 ± 0.04 | 24.5 ± 0.17 |
| *phm>CG30054*-RNAi | developmental delay (Yamanaka et al., 2015) | 7 | 0.25 ± 0.03 | 24.8 ± 0.78 |
| *phm>CG17760*-RNAi | Gαq protein (Yamanaka et al., 2015) | 5 | 0.27 ± 0.04 | 24.2 ± 0.88 |
| *phm>egfr* | precocious pupariation (Cruz et al., 2020) | 4 | Eclosion failure | Eclosion failure |
| *phm>pten*-RNAi | PG growth (Mirth et al., 2005) | 5 | 0.24 ± 0.02 | 24.0 ± 0.36 |
| *phm>pi3k*-RNAi | 20-hydroxiecdysone titers (Caldwell et al., 2005; Colombani et al., 2005) | 5 | 0.31 ± 0.03 | 24.4 ± 0.20 |
| *phm>InR*-RNAi | ecdysone biosynthesis (Colombani et al., 2005) | 3 | 0.25 ± 0.05 | 24.1 ± 0.21 |
| *dilp2-3, 5* null | nutritional input to the PG (Kannangara et al., 2021) | 2 (1621) | 0.16 ± 0.04 | 24.0 ± 0.44 |
| *dilp 7* null | insulin signaling (Kannangara et al., 2021) | 3 (2231) | 0.32 ± 0.03 | 24.0 ± 0.16 |
| *dilp 8* null | ecdysone biosynthesis (Garelli et al., 2012) | 2 (1023) | 0.39 ± 0.04 | 24.5 ± 0.25 |
| *lgr3* null | developmental delay (Colombani et al., 2015) | 3 (1145) | 0.40 ± 0.04 | 23.5 ± 0.28 |
| *lsp2>Δ901* | fat body clock (Xu et al., 2008) | 5 | 0.36 ± 0.03 | 24.3 ± 0.26 |

Periods were calculated using Autocorrelation analysis. (N Total number of different experiment performed. (n) indicates the number of flies emerged out from the pupal case; (SEM) standard error; (RI) rhythmicity index. Records are considered arrhythmic if RI<0.1.

#### Table S2. Sequence of PCR Primers used.

| Oligonucleotides | Sequence of primer pair |
|---|---|
| *torso* forward primer | CCAGTGATCTCTTGCAGCTAC |
| *torso* reverse primer | AGTCTGTGTTTAAGGGCGG |
| *rp49* forward primer | TGTGATGGGAATTCGTGGG |
| *rp49* reverse primer | ATCTTGGGCCTGTATGCTG |

### Table S3. Primary antibodies used

| Primary antibody | Procedure | Source | Dilution | Catalogue#/ Reference |
|---|---|---|---|---|
| Rat α-mCherry monoclonal 16D7 | Double staining, BAcTrace | Invitrogen | 1:1000 | M11217 |
| Rabbit α-GFP polyclonal | Double staining, BAcTrace, syb-GRASP, CaLexA | Chromotek | 1:1000 | PABG1 |
| Guinea pig α-RFP polyclonal | *trans-Tango* MkII | Meet Zandawala | 1:10000 | |
| Rabbit α-PTTH-polyclonal | Double staining, *trans-Tango* MkII, CaLexA | this paper | 1:10000 | this paper |
| Guinea pig, α-PTTH polyclonal | Immunocytochemistry | Michael O’Connor | 1:500 | (McBrayer et al., 2007) |
| Rabbit α-GFP polyclonal | Immunocytochemistry | Thermofisher | 1:1000 | A11122 |
